# An expanded toolbox for genetic access to molecularly defined cortical intratelencephalic neuron subtypes

**DOI:** 10.64898/2026.10.30.762849

**Authors:** Jeffrey B. Russ, Jessica S. Zhang, X. Hermione Xu, Weixin Zhong, Teresa Liang, Zixuan Qiu, Sanjeev Kaushalya, Yida Zhang, Peregrine Ke-Lind, Z. Josh Huang

**Author notes:** Corresponding authors: Jeffrey Russ, 311 Research Drive, Durham, NC 27710, Z. Josh Huang, 311 Research Drive, Durham, NC 27710. These authors contributed equally to this work (Jeffrey B. Russ, Jessica S. Zhang).

## Abstract

Among the multiple cortical glutamatergic neuron classes, intratelencephalic (IT) neurons represent the largest class and project within the cortex and striatum. IT neurons have been amplified and diversified throughout mammalian evolution and are thought to underlie increasingly complex mammalian cognition. However, despite their diversity and significance in cortical processes, few existing tools connect the molecular signatures to the projection patterns of distinct IT subtypes, limiting our ability to understand their organization and function at a subtype resolution. Here, we illustrate six novel gene knock-in mouse lines based on IT-specific markers, labeling subpopulations with unique cortico-cortical and corticostriatal projection patterns. These include two broad subpopulations with upper-layer (*Cux1-Flp*) versus lower-layer (*Kcnh5-CreER*) expression biases, two narrow upper-layer subpopulations with contrasting callosal (*Wfs1-Flp*) and local (*Igfn1-Flp*) projection patterns, and two narrow lower-layer subpopulations with contrasting bilateral (*Deptor-CreER*) versus ipsilateral-predominant (*Ptprm-CreER*) corticostriatal projection patterns. Upper-layer *Wfs1*^+^ and lower-layer *Ptprm*^+^ IT neurons also project to perirhinal/ectorhinal cortex with densities that vary across cortical areas. Finally, we use *in vivo* calcium imaging in mice performing a reach-withdraw-drink task to demonstrate subtype-specific temporal dynamics in upper-layer *Wfs1*^+^ versus lower-layer *Deptor*^+^ neurons. Our work expands the toolbox of transgenic mice to access molecularly defined IT neuron subpopulations.

**Impact Statement:** Six transgenic mouse lines provide unprecedented genetic access to molecularly defined upper- and lower-layer cortical intratelencephalic (IT) neuron subtypes, which demonstrate distinct axonal projections patterns and activity during a behavioral task.

## INTRODUCTION

The telencephalon is the rostral-most portion of the mammalian central nervous system, encompassing the cerebral cortex, basal ganglia, and limbic system. Glutamatergic excitatory projection neurons of the cerebral cortex can be divided into two main classes based on whether they project within or away from the telencephalon (Fame et al., 2011; Greig et al., 2013; Harris & Shepherd, 2015). Intratelencephalic (IT) neurons make intrahemispheric and callosal cortico-cortical connections, as well as ipsilateral and/or contralateral corticostriatal projections (Fame et al., 2011). Extratelencephalic (ET) neurons, in contrast, comprise pyramidal tract (PT), corticothalamic (CT), and other corticofugal projection neurons (CFuPN), which are the major output neurons of the cortex and project to targets such as the thalamus, tectum, pons, or spinal cord (Greig et al., 2013; Harris & Shepherd, 2015).

Throughout vertebrate – particularly mammalian – evolution, a shift toward indirect neurogenesis, wherein neurons arise from intermediate cortical progenitors instead of directly from radial glia, served to amplify and diversify IT neurons more than other cortical excitatory neuron classes (Cardenas & Borrell, 2020; Huilgol et al., 2023). The enhanced number and density of IT neurons, especially in upper layers of primate and human cortex, is thought to underpin the increasingly complex cognitive capacities of higher-order mammals. Conversely, IT neuron dysfunction is also postulated to be uniquely perturbed in cognitive neurodevelopmental disorders, such as autism (Starr & Fraser, 2025; Velmeshev et al., 2019; Wang et al., 2025).

Despite their well-documented heterogeneity, there remains a limited understanding of discrete IT neuron subtypes defined by their molecular signatures and hodological patterns. Classic IT markers, such as *Satb2*, *Cux1*, and *Plxnd1* broadly label IT neurons across laminae (Alcamo et al., 2008; Britanova et al., 2008; Matho et al., 2021; Nieto et al., 2004). Recent advances in single-cell profiling have delineated upper- and lower-layer subtypes at higher transcriptomic resolution in mouse and human cortex (Berg et al., 2021; Cheng et al., 2022; Fang et al., 2022), but do not connect gene expression to axonal projection patterns. Electrophysiological and hodological studies provide a general overview of basic IT projection motifs, though not at the level of molecularly defined IT subtypes (Gao et al., 2022; Shepherd, 2013; Ueta et al., 2013; Ueta et al., 2019; Zingg et al., 2014). Studies that delineate projection patterns of molecularly defined IT subtypes rely on a limited number of transgenic mouse lines that often label broad inter-laminar populations (Liu et al., 2024; Munoz-Castaneda et al., 2021; Peng et al., 2021).

In this study, we showcase a toolbox of novel gene knock-in mouse lines that leverage more specific IT marker genes to drive Flp or CreER recombinase expression, providing genetic access to distinct IT subtypes. We identify two upper-layer populations (*Wfs*^+^, *Igfn1*^+^), two lower-layer populations (*Deptor*^+^, *Ptprm*^+^), and a broad marker that excludes the superficial-most neurons of layer 2 (*Kcnh5^+^*). For comparison, we also capitalize on the well-characterized IT marker *Cux1* to generate a *Cux1-Flp* line that constitutively marks all cells with a history of *Cux1* expression. For each marker-defined subtype, we map cell distribution across the brain via serial two-photon tomography (STPT) and perform anterograde viral tracing to characterize axonal projection patterns from primary motor cortex (MOp) and primary sensory cortex (SSp). We also identify that upper-layer *Wfs1*^+^ and lower-layer *Ptprm*^+^ IT neurons both show prominent projections to perirhinal/ectorhinal cortex (PERI/ECT), a critical point of convergence from across cortical areas with input to the medial prefrontal cortex (mPFC), lateral entorhinal cortex (ENTl), and hippocampal formation. We compare differences in the cortical areas giving rise to PERI/ECT input for these two IT populations. Finally, we demonstrate contrasting temporal dynamics between upper-layer *Wfs1*^+^ and lower-layer *Deptor*^+^ IT neurons via two-photon calcium imaging during an in vivo reach-withdraw-drink (RWD) task. Together, these novel mouse lines illuminate distinct differences in laminar position, projection patterns, and activity across an expanded variety of molecularly defined IT subtypes.

## RESULTS

### IT marker identification and validation

To identify candidate genes with expression restricted to different subsets of IT neurons, we reviewed literature (Kim et al., 2020), single-cell transcriptomic atlases (Cheng et al., 2022; Yao et al., 2023), and the Allen adult mouse cortex *in situ* hybridization (ISH) database (Lein et al., 2007; https://mouse.brain-map.org; Supplementary Figure 1A). We then curated a list of top candidate markers that appeared to have expression biases in IT subsets, including *Wfs1* and *Igfn1* in upper cortical layers, *Kcnh5* in middle and lower cortical layers, and *Deptor* and *Ptprm* in lower cortical layers. *Cux1* is a well-characterized marker of IT neurons in layers 2-4 and was included for comparison.

We validated the IT-enriched expression of our selected candidates through several means: 1) co-expression with the pan-IT marker *Satb2* (Alcamo et al., 2008; Britanova et al., 2008) via hybridization chain reaction (HCR) ISH (Figure 1E,F), co-expression with *Satb2* via multiplexed error-robust fluorescence ISH (MERFISH; Supplementary Figure 2), and expression in annotated IT subtypes in a single-cell transcriptomic mouse cortex atlas (Yao et al., 2023; Supplementary Figure 1B,C). Via HCR ISH, *Wfs1*, *Igfn1*, *Kcnh5*, and *Deptor* all showed near complete overlap (>97.6 ± 2.1%) with *Satb2* in mouse cortex (Figure 1E). In cortical excitatory projection neurons, *Ptprm* expression is restricted to layer 5 IT neurons, although it is also expressed in inhibitory interneurons and endothelial cells throughout all cortical layers (Yao et al., 2023). When analysis of *Satb2* co-expression was limited to cells in layer 5, *Ptprm* co-labeled with *Satb2* in 75.1 ± 4.8% of layer 5 cells (Figure 1E).

**Figure 1:**
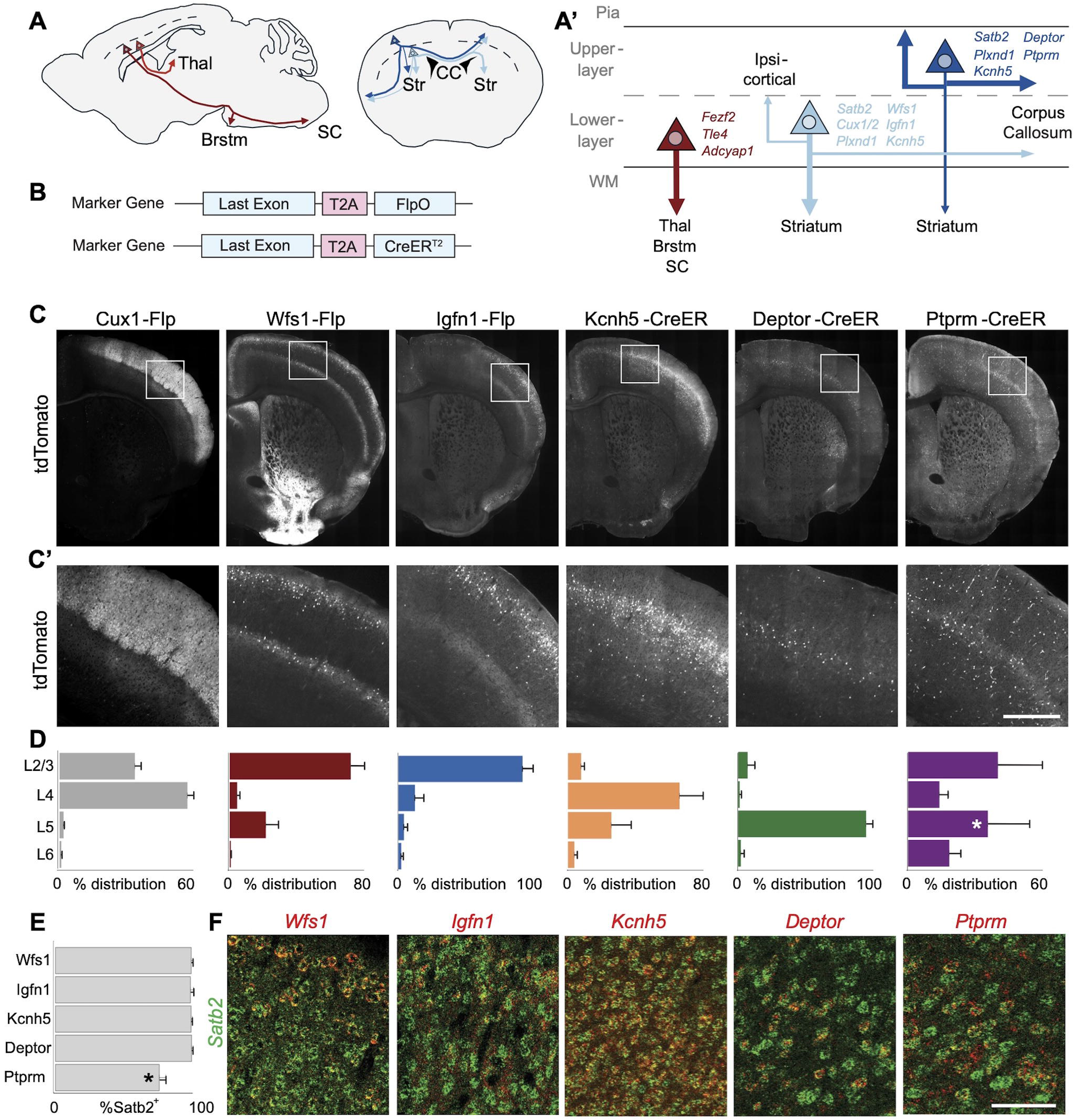
Six novel knock-in mouse lines provide genetic access to distinct IT neuron subtypes. **(A)** Extratelencephalic (ET) excitatory projection neurons (left, red) reside in lower cortical layers (dashed line) and project out of the cortex to the thalamus, brainstem, and spinal cord. Intratelencephalic (IT) neurons (right, blue) reside in upper and lower cortical layers and make callosal, corticostriatal, and intrahemispheric connections. **(A’)** Upper layer IT neurons (right, dark blue) classically make preferential callosal and intrahemispheric cortico-cortical connections, while lower layer IT neurons (middle, light blue) classically make preferential corticostriatal connections. Representative genetic markers for each cell type are listed. **(B)** Schematic map of knock-in mouse line generation. A self-cleaving T2A followed by Flpo or CreER^T2^ was inserted after the last exon of each IT marker of interest. **(C)** Coronal sections from whole brain of six Flp- or CreER-dependent mouse lines crossed with Flp-dependent (Ai65F) or Cre-depedent (Ai14) tdTomato fluorescent reporters and imaged via STPT. **(C’)** Coronal images of SSp for each reporter-crossed IT line from the boxed regions in (C). **(D)** Fluorescent reporter-labeled cell body density across cortical laminae for each IT line. Mean count per layer across N=2 coronal cortical sections from each of N≥3 mice per line is plotted. Error bars indicate standard deviation. Asterisk in layer 5 for *Ptprm-CreER* indicates where *Ptprm*^+^ excitatory projection neurons reside. **(E)** Percent overlap of each IT marker with the pan-IT marker *Satb2* through HCR ISH in (F). Plots represent the mean percent overlap of each IT marker with *Satb2* for N=2 coronal cortical sections from N=3 mice. Error bars indicate standard deviation. Asterisk for *Ptprm* indicates that analysis of *Ptprm*/*Satb2* overlap was restricted to layer 5, where *Ptprm*^+^ excitatory projection neurons reside. **(F)** Representative HCR ISH images of each IT marker co-expression with *Satb2*. Scalebar in (C) = 1mm, (C’) = 500µm, (F) = 100 µm.

Using the existing IT subtype annotations from the Allen Institute in the mouse cortex transcriptomic atlas (Yao et al., 2023), we found that 85.7% of *Cux1*^+^ excitatory neurons were in the IT class (Supplementary Figure 1C), primarily in layers 2/3 (32.9%) and 4/5 (50.1%), consistent with its known upper-layer IT expression bias (Matho et al., 2021; Nieto et al., 2004). Similarly, 88.8% of *Kcnh5*^+^ excitatory neurons were of the IT class (Supplementary Figure 1C), though with a medium- and lower-layer bias (68.9% characterized as layer 4/5 IT and 10.8% as layer 5 IT). 92.9% of *Wfs1*^+^ excitatory neurons were characterized in the IT class with 69.7% as layer 2/3 IT and 13.0% as layer 4/5 IT. 99.8% of *Igfn1*^+^ excitatory neurons were in the IT class, with 33.2% labeled as layer 2/3 IT and 52.8% labeled as layer 4/5 IT. For *Deptor*, 80% were characterized in the IT class with 31.3% annotated as layer 4/5 IT and 41.5% annotated as layer 5 IT. The most common non-IT annotation for *Deptor*^+^ cells was layer 6 corticothalamic (CT; 15.9%). Finally, 97.2% of *Ptprm*^+^ excitatory neurons were of the IT class, with 55.7% annotated as layer 4/5 IT and 41.1% annotated as layer 5 IT.

### Cux1-Flp

*Cux1* is a well-described transcription factor expressed early in corticogenesis. *Cux1* mRNA is first detected in the ventricular zone as early as E14 with increasing Cux1 immunoreactivity in post-mitotic neurons destined for upper cortical layers by E16 (Ferrere et al., 2006; Nieto et al., 2004). As the cortex matures, *Cux1* expression is restricted to excitatory neurons in layers 2 through 4 with very little expression in deeper layers, though lower levels of scattered *Cux1* expression can be observed in some interneuron subpopulations (Yao et al., 2023). Given its spatial restriction to upper cortical layers, where most IT neurons reside, *Cux1* is typically considered a “classic” upper-layer IT marker and is thought to play a role in dendrite morphology (Cubelos et al., 2015) and activity-dependent callosal axon crossing (Rodriguez-Tornos et al., 2016).

Our lab previously generated a knock-in *Cux1-CreER* line (Matho et al., 2021) that allows for temporally controlled labeling of *Cux1*^+^ neurons, but to date no transgenic mouse line exists for constitutive labeling of *Cux1*^+^ neurons. We therefore generated a *Cux1-Flp* knock-in mouse line that introduces P2A-Flpo between the final exon (exon 24) and the 3’ UTR of the endogenous *Cux1* gene. Crossing our *Cux1-Flp* line with the Flp-dependent tdTomato-reporter line *Ai65F* (Daigle et al., 2018) demonstrated the expected endogenous pattern of dense cell soma labeling in layers 2 through 4 of the cerebral cortex, with highest density in SSp. Dense *Cux1*^+^ cell body labeling was also observed in the main olfactory bulb, anterior olfactory nucleus, and claustrum (Supplementary Figure 3A,B).

Anterograde labeling from MOp using a Flp-dependent EYFP packaged in an AAV2/9 capsid revealed dense cortico-cortical projections to ipsilateral SSp and secondary sensory cortex (SSs) with sparser projections to ipsilateral secondary motor cortex (MOs; Figure 2A,B). Dense callosal axons were also observed projecting to contralateral homotopic MOp, as well as sparser heterotopic projections to contralateral MOs, SSp, and SSs. Across these cortical areas, axons were observed in all cortical layers, but with the highest densities in layers 2/3 and 5. Bilateral cortico-cortical projections were also observed in the dense area of lateral cortex that includes PERI/ECT, with similar sparser projections to bilateral insular cortex (INS) more anteriorly. Finally, corticostriatal projections were observed primarily in central and dorsolateral regions of the ipsilateral striatum (Figure 2A,B).

**Figure 2:**
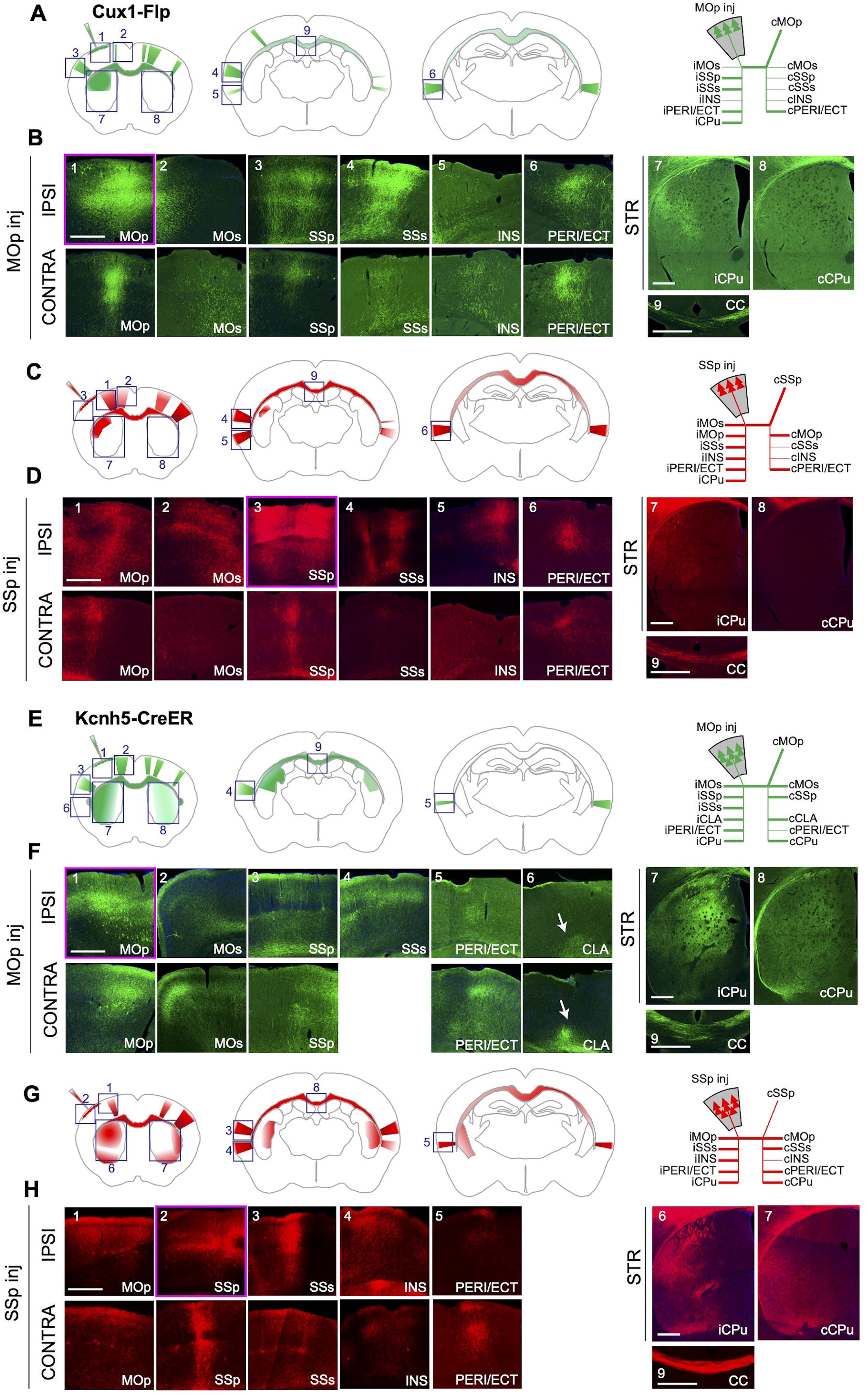
*Cux1-Flp* and *Kcnh5-CreER* capture broad but distinct IT neuron populations. **(A,B)** EYFP-labeled anterograde projections from MOp of *Cux1-Flp* mice. **(A)** Schematic illustration (left) of major *Cux1*^+^ axonal projections from MOp in coronal cross sections of mouse brain. Stick diagram (right) of major (thick lines) and minor (thin lines) ipsilateral and contralateral *Cux1*^+^ axonal projections from MOp. **(B)** High-magnification confocal images of ipsilateral and contralateral axonal projections from MOp (magenta outline) to each of the numbered boxed regions in (A). **(C,D)** mCherry-labeled anterograde projections from SSp of *Cux1-Flp* mice. **(C)** Schematic illustration (left) of major *Cux1*^+^ axonal projections from SSp. Stick diagram (right) of major (thick lines) and minor (thin lines) ipsilateral and contralateral *Cux1*^+^ axonal projections from SSp. **(D)** High-magnification confocal images of ipsilateral and contralateral axonal projections from SSp (magenta outline) to each of the numbered boxed regions in (C). **(E,F)** EGFP-labeled anterograde projections from MOp of *Kcnh5-CreER* mice. **(E)** Schematic illustration (left) of major *Kcnh5*^+^ axonal projections from MOp. Stick diagram (right) of major (thick lines) and minor (thin lines) ipsilateral and contralateral *Kcnh5*^+^ axonal projections from MOp. **(F)** High-magnification confocal images of ipsilateral and contralateral axonal projections from MOp (magenta outline) to each of the numbered boxed regions in (E). **(G,H)** tdTomato-labeled anterograde projections from SSp of *Kcnh5-CreER* mice. **(G)** Schematic illustration (left) of major *Kcnh5*^+^ axonal projections from SSp. Stick diagram (right) of major (thick lines) and minor (thin lines) ipsilateral and contralateral *Kcnh5*^+^ axonal projections from SSp. **(H)** High-magnification confocal images of ipsilateral and contralateral axonal projections from MOp (magenta outline) to each of the numbered boxed regions in (G). All anterograde projections were confirmed through observation across N=3 mice per line. IPSI = ipsilateral, CONTRA = contralateral, i = ipsilateral, c = contralateral, inj = injection, MOp = primary motor cortex, MOs = secondary motor cortex, SSp = primary somatosensory cortex, SSs = secondary somatosensory cortex, INS = insular cortex, PERI/ECT = perirhinal/ectorhinal cortex, CLA = claustrum, CPu = caudate/putamen, CC = corpus callosum. Scalebars = 500 µm.

Anterograde labeling from SSp using a Flp-dependent mCherry delivered via AAV2/9 revealed a similar projection pattern, with dense ipsilateral labeling in MOp and MOs, as well as SSs (Figure 2C,D). Dense callosal axons were observed with homotopic projections to contralateral SSp, as well as heterotopic projections to contralateral MOp and sparser projections to SSs. While axonal projections were observed across cortical layers in both ipsilateral and contralateral cortices, axonal density was highest in layers 2/3 and 5. Prominent bilateral projections from SSp were again observed in ipsilateral and contralateral PERI/ECT, and sparser ipsilateral-predominant projections to INS. Corticostriatal projections from *Cux1*^+^ neurons in SSp were largely restricted to the ipsilateral striatum with central and dorsolateral distributions (Figure 2C,D).

### Kcnh5-CreER

*Kcnh5* encodes a voltage-dependent potassium channel subunit (Saganich et al., 1999). *Kcnh5* is excluded from the most superficial cortical layers, and its densest expression is in layer 4. Thus, prior studies have utilized *Kcnh5* as an evolutionarily conserved marker of neurons receiving thalamic input (Briscoe et al., 2018; Dugas-Ford et al., 2012). However, *Kcnh5* is not exclusively a layer 4 marker. *Kcnh5* expression is excluded from superficial layer 2 and expressed largely in IT neurons that extend as deep as layer 6 (Lein et al., 2007; https://mouse.brain-map.org; Figure 1C’; Supplementary Figure 1A). In primary visual cortex (VIS), the expression of *Kcnh5* can distinguish IT neurons that reside below layer 2 (Cheng et al., 2022) and project to anterolateral rather than posteromedial higher order visual areas (Kim et al., 2020).

Given the contrast in laminar expression with more superficial *Cux1*^+^ IT neurons, we generated a transgenic *Kcnh5-CreER* knock-in mouse line. This line introduces T2A-IRES-CreER^T2^ after the final exon (exon 11) and the 3’ UTR of the endogenous *Kcnh5* gene. When crossed with the Cre-dependent tdTomato reporter line *Ai14* (Madisen et al., 2010), *Kcnh5*^+^ cell bodies were observed in the expected cortical laminar distribution, particularly in deep layer 2/3, layer 4, layer 5, and scattered cells in layer 6 (Figure 1C-D). Subcortically, *Kcnh5*^+^ cell bodies were also observed in olfactory structures (main olfactory bulb, anterior olfactory nucleus, and olfactory tubercle), piriform cortex, the lateral and intercalated amygdalar nuclei, the bed nucleus of the anterior commisure, the presubiculum, and the postsubiculum (Supplementary Figure 4A,B).

Anterograde ipsilateral projections of *Kcnh5*^+^ IT neurons from MOp, labeled via Cre-dependent EGFP, were observed in SSp and SSs with dense callosal projections to contralateral homotopic MOp and MOs, as well as dense heterotopic projections to contralateral SSp (Figure 2E,F). Projections to bilateral PERI/ECT with a prominent contralateral projection and minor ipsilateral projection were also observed. Axonal projections to these cortical areas ended primarily in layers 1, 2/3, and 5. Unique to the *Kcnh5-CreER* line, we observed bilateral projections to the claustrum (Figure 2E,F), with the contralateral claustrum more densely labeled than the ipsilateral one, suggesting that *Kcnh5* labeling captures a unique cortico-claustrum subpopulation. Bilateral corticostriatal projections were also prominently observed, with a relatively diffuse projection pattern that spanned the dorsoventral axis of both striata (Figure 2E,F).

Anterograde labeling from SSp via a Cre-dependent tdTomato reporter demonstrated ipsilateral projections to MOp and SSs (Figure 2G,H). Dense callosal axons project primarily to homotopic SSp and heterotopic contralateral MOp and SSs. Axon terminations spanned cortical laminae with the highest density in layer 1, layer 2/3, and upper layer 5. In lateral cortex, projections from SSp were also observed in bilateral INS and PERI/ECT. *Kcnh5*^+^ IT neurons in SSp also sent bilateral corticostriatal projections that favored the ipsilateral striatum and exhibited a broad distribution spanning its dorsoventral axis. Ipsilateral corticostriatal projections also exhibited a greater posterior extension compared to the contralateral striatum. Finally, sparse tdTomato-labeled axons were noted passing through the ipsilateral striatum, internal capsule, and cerebral peduncle suggesting that a minority of *Kcnh5*^+^ cortical neurons are PT neurons from the ET class.

### Wfs1-Flp

*Wfs1* encodes Wolframin, an endoplasmic reticulum glycoprotein with roles in regulating calcium homeostasis and endoplasmic reticulum stress (Fonseca et al., 2010; Takei et al., 2006). In the mouse cortex, *Wfs1* expression is restricted to the superficial-most cells of cortical layer 2/3 (Lein et al., 2007; Yao et al., 2023; Zariwala et al., 2011; Figure 1C-D, Supplementary Figure 1A). *Wfs1* expression is detected in the embryonic striatum as early as E15.5, and expression in the superficial layers of lateral and ventral cortical areas, such as PERI, INS, and piriform cortex, begins around E18.5 (Tekko et al., 2014). In dorsal isocortex, beginning around P0, *Wfs1* expression begins first in layer 5, starting in medial regions, including the cingulate, retrosplenial, and motor cortex, and then progressing across the cortex. Expression in layer 5 is transient, however, and the sustained expression of *Wfs1* across layer 2/3 of the isocortex begins at P4 (Tekko et al., 2014). In adult isocortex, *Wfs1* expression remains restricted to neuronal somata in layers 2/3, though detectable Wfs1 protein in layer 5 is thought to reflect axonal projections from superficial *Wfs1*^+^ neurons to deeper cortical layers (Tekko et al., 2017).

Previously, a bacterial artificial chromosome (BAC)-transgenic *Wfs1-CreER* line has been generated (Madisen et al., 2010), though no constitutive recombinase line exists to label *Wfs1*^+^ neurons. We therefore generated a constitutive *Wfs1-Flp* knock-in mouse line that incorporates T2A-Flpo into the 3’ UTR of the endogenous *Wfs1* gene following the final exon (exon 8). Crossing our *Wfs1-Flp* line to the Flp-dependent *Ai65F* reporter line (Daigle et al., 2018) recapitulates endogenous patterns of expression across the adult mouse brain, with dense labeling in expected regions, such as the nucleus accumbens, CA1 of the hippocampus, and the olfactory tubercles, among other regions (Supplementary Figure 3C,D). In cortex, *Wfs1*^+^ somata were primarily located in superficial layer 2/3, with sparser labeling in layer 5, reflecting the developmental history of transient layer 5 expression (Tekko et al., 2014). *Wfs1*^+^ neurites were also dense throughout layer 5, likely reflecting a combination of local neurites from *Wfs1*^+^ layer 5 neurons and projections from layer 2/3 *Wfs*^+^ neurons (Tekko et al., 2017).

Through Flp-dependent anterograde tracing, we observed that *Wfs1*^+^ neurons in MOp were largely restricted to superficial layer 2/3 with ipsilateral projections to layers 2/3 and 5 of SSp and SSs (Figure 3A,B). Viral tracer was also observed in ipsilateral MOs but appeared to be a mixture of both axonal projections and cell body labeling, which may reflect both local diffusion of viral tracer and close interconnectivity of MOp and MOs. Dense callosal projections to layers 2/3 and 5 of contralateral homotopic MOp and MOs, as well as heterotopic projections to layers 2/3 and 5 of contralateral SSp were also observed. Sparse heterotopic projections in contralateral SSs were also present. EYFP projections from *Wfs1*^+^ neurons in MOp were also found densely concentrated in bilateral PERI/ECT, mirroring a projection motif that has been described for a limited population of *Wfs1*^+^ neurons in rat MOs (Ueta et al., 2019). Sparser projections to bilateral INS just anterior to PERI/ECT were also observed. *Wfs1*^+^ corticostriatal projections from MOp were found concentrated in the ipsilateral striatum, largely restricted to a lateral striatal territory that overlaps with previously described somatotopic projections from mouse sensorimotor cortex (Hintiryan et al., 2016; Wang et al., 2026). Scant contralateral corticostriatal signal was highly localized to dorsal striatum, which we hypothesize based on canonical projection motifs (Shepherd, 2013) are more likely to arise from the minor population of layer 5 *Wfs1*^+^ neurons.

**Figure 3:**
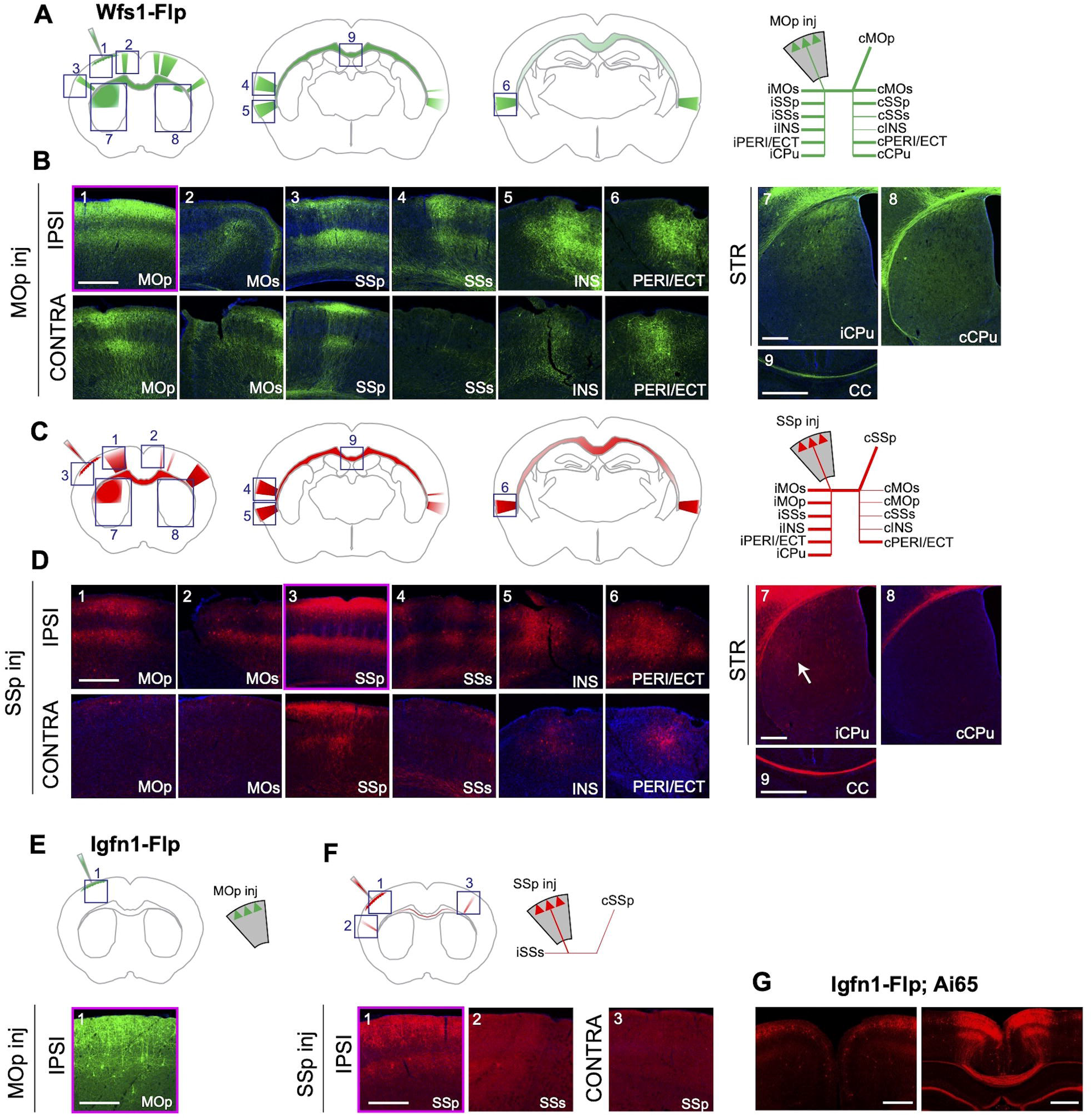
*Wfs1-Flp* and *Igfn1-Flp* label distinct upper-layer IT neurons with contrasting long-range projection patterns. **(A,B)** EYFP-labeled anterograde projections from MOp of *Wfs1-Flp* mice. **(A)** Schematic illustration (left) of major *Wfs1*^+^ axonal projections from MOp. Stick diagram (right) of major (thick lines) and minor (thin lines) ipsilateral and contralateral *Wfs1*^+^ axonal projections from MOp. **(B)** High-magnification confocal images of the boxed regions in (A), demonstrating EYFP-positive anterograde projections from MOp (magenta outline) of *Wfs1-Flp* mice to ipsilateral MOs, SSp, SSs, INS, and PERI/ECT. Contralateral *Wfs1*^+^ projections from MOp are observed in homotopic MOp and heterotopic MOs, SSp, INS, and PERI/ECT with almost no projections to contralateral SSs. **(C,D)** mCherry-labeled anterograde projections from SSp of *Wfs1-Flp* mice. **(C)** Schematic illustration (left) of major *Wfs1*^+^ axonal projections from SSp. Stick diagram (right) of major (thick lines) and minor (thin lines) ipsilateral and contralateral *Wfs1*^+^ axonal projections from SSp. **(D)** High-magnification confocal images of the boxed regions in (C), demonstrating mCherry-positive anterograde projections from SSp (magenta outline) of *Wfs1-Flp* mice in ipsilateral MOp, MOs, SSs, INS, and PERI/ECT. Dense homotopic projections are observed in contralateral SSp, sparser projections in contralateral SSs, INS, and PERI/ECT, and almost no projections to contralateral MOp/MOs. **(E)** Schematic illustration (top left) and stick diagram (top right) of *Igfn1*^+^ axonal projections from MOp in coronal cross sections of mouse brain. EYFP-positive anterograde labeling from MOp (bottom, magenta outline) of *Igfn1-Flp* mice showed no long-range projections. **(F)** Schematic illustration (top left) and stick diagram (top right) of *Igfn1*^+^ axonal projections from SSp in coronal cross sections of mouse brain. mCherry-positive anterograde projections from SSp (bottom, magenta outline) of *Igfn1-Flp* mice shows scant labeling in ipsilateral SSs and contralateral homotopic SSp. **(G)** Anterior coronal section (left) from an *Igfn1-Flp;Ai65F* mouse showing that despite tdTomato-positive cell bodies in bilateral sensorimotor cortex, there are nearly no callosal axons observed. More posterior (right), the only prominent tdTomato-labeled callosal axons in *Igfn1-Flp* mice arise from retrosplenial cortex and cross in the dorsal corpus callosum, consistent with callosal axon topography from medial cortical neurons (Poulopoulos et al., 2024). All anterograde projections were confirmed through observation across N=3 mice per line. IPSI = ipsilateral, CONTRA = contralateral, i = ipsilateral, c = contralateral, inj = injection, MOp = primary motor cortex, MOs = secondary motor cortex, SSp = primary somatosensory cortex, SSs = secondary somatosensory cortex, INS = insular cortex, PERI/ECT = perirhinal/ectorhinal cortex, CPu = caudate/putamen, CC = corpus callosum. Scalebars = 500 µm.

*Wfs1*^+^ neurons in SSp were also largely restricted to superficial layer 2/3 with dense ipsilateral projections to SSs. Ipsilateral MOp and MOs contained both axons and cell bodies, likely reflecting a mixture of primary projections with occasional retrograde uptake of AAV2/9 (Castle et al., 2014) by MOp/MOs IT neurons with extensive interconnectivity with ipsilateral SSp. *Wfs1*^+^ neurons in SSp also exhibited dense homotopic callosal projections to layers 2/3 and 5 of contralateral SSp (Figure 3C,D). Sparse contralateral heterotopic projections were seen in SSs, as well as very sparse heterotopic projections in contralateral MOp/MOs. As with projections from motor cortex, *Wfs1*^+^ neurons in primary sensory cortex projected densely to bilateral PERI/ECT with ipsilateral more than contralateral INS projections just anterior to PERI/ECT. Corticostriatal projections from SSp were solely ipsilateral and restricted to a similar dorsolateral region of the striatum as projections from MOp.

### Igfn1-Flp

*Igfn1* encodes “immunoglobulin-like and fibronectin type III domain containing 1,” a cytosolic protein enriched in skeletal muscle that interacts with eukaryotic translation elongation factors (Mansilla et al., 2008). In cortex, *Igfn1* is expressed after P14 in a subset of IT neurons that primarily reside in deeper layers 2/3 (Cheng et al., 2022). Cheng et al. (2022) demonstrated that, at least in visual cortex, both *Igfn1* expression and transcriptional maturation of the *Igfn1*^+^ subset depend on early visual stimulation.

Given its unique and narrow expression pattern in a subset of layer 2/3 IT neurons (Supplementary Figures 1 and 2), we generated *Igfn1-Flp* mice, which constitutively express T2A-Flpo between the final exon of *Igfn1*, exon 27, and the 3’ UTR. Breeding this line with the Flp-dependent reporter line *Ai65F* demonstrates expected endogenous labeling patterns in superficial layers of mouse cortex (Figure 1C-D), including expression that does not extend fully to the superficial-most boundary of layer 2/3. Across whole brain, *Igfn1*^+^ cell bodies are also apparent in retrosplenial (RSP) and superficial piriform cortex (Supplementary Figure 3E,F). Within the isocortex, *Igfn1* demonstrates stark differential expression across cortical areas with enriched cell body density in SSp and SSs, VIS, and RSP (Figures 1C, 3G and Supplementary Figure 3E,F).

To delineate the projection patterns of *Igfn1*^+^ cortical neurons, we employed anterograde AAV2/9 tracing with Flp-dependent EYFP or mCherry from MOp and SSp, respectively (Figure 3E,F). AAV2/9 labeled superficial cortical neurons in layer 2/3 with only scant long-range projections. Particularly in SSp, dense axonal projections were apparent in layer 5, recapitulating a well-known motif of interconnectivity between layer 2/3 and layer 5 (Harris & Shepherd, 2015). Sparse *Igfn1*^+^ projections from SSp to ipsilateral SSs, as well as sparse callosal axons to homotopic contralateral SSp, were also observed. Indeed, when crossed with *Ai65F* reporter mice, *Igfn1*^+^ callosal axons are also sparse, appearing to derive primarily from RSP, with a minor more ventral component from sensory cortex (Figure 3G). Likewise, *Igfn1*^+^ neurons did not appear to extend corticostriatal projections from either MOp or SSp. Given the dearth of long-range anterograde projections from *Igfn1*^+^ neurons, we infer that these may represent a unique population of local or intrahemispheric IT neurons.

### Deptor-CreER

In addition to our upper-layer IT lines, we likewise generated knock-in mouse lines for lower-layer IT markers. *Deptor*, which stands for “Disheveled, Egl-10, and Plekstrin (DEP) domain-containing mechanistic target of rapamycin (mTOR)-interacting protein,” is an mTOR binding partner that inhibits the kinase activity of mTOR (Vega et al., 2022). In cortex, *Deptor* is expressed in a narrow band of layer 5 neurons (Figures 1C-D; Supplementary Figures 1 and 2; Cheng et al., 2022). To explore cortical *Deptor* expression and axonal projection patterns of *Deptor*^+^ IT neurons, we generated transgenic knock-in CreER mice, wherein T2A-IRES-CreER^T2^ was inserted between exon 9 of the endogenous *Deptor* gene and its 3’ UTR.

Mice were crossed with the Cre-dependent *Ai14* reporter (Madisen et al., 2010) and induced with tamoxifen. In isocortex, *Deptor*^+^ neurons were restricted to the expected narrow band in layer 5 (Figure 1C’). Subcortically, *Deptor*^+^ cell bodies are also densely clustered in a small ventrolateral area of the striatum, as well as subsets of the subiculum, hypothalamus, amygdala, and pontine gray matter (Supplementary Figure 4C,D). Choroid plexus also demonstrates strong *Deptor* expression (Supplementary Figure 4D).

Anterograde AAV2/9 viral tracing from MOp or SSp via Cre-dependent EGFP or tdTomato, respectively, demonstrated distinct area-specific *Deptor*^+^ projection patterns (Figure 4A-D). EGFP-labeled *Deptor*^+^ IT neurons from MOp exhibited scant projections to layer 2/3 of ipsilateral SSs. *Deptor*^+^ IT neurons in MOp also make sparse callosal projections to layers 2/3 and 5 of homotopic contralateral MOp and heterotopic SSp with no other notable contralateral cortico-cortical projections. *Deptor*^+^ IT neurons in MOp also demonstrated a unique corticostriatal projection pattern. In anterior striatum, *Deptor*^+^ IT neurons from MOp exhibited symmetric bilateral corticostriatal projections that spanned the dorsoventral axis of the striata. However, just posterior in the central striatum, corticostriatal projections from motor cortex exhibited asymmetric projections primarily to ipsilateral, dorsolateral striatum with minor projections to the very ventral edge of contralateral striatum (Figure 4B).

**Figure 4:**
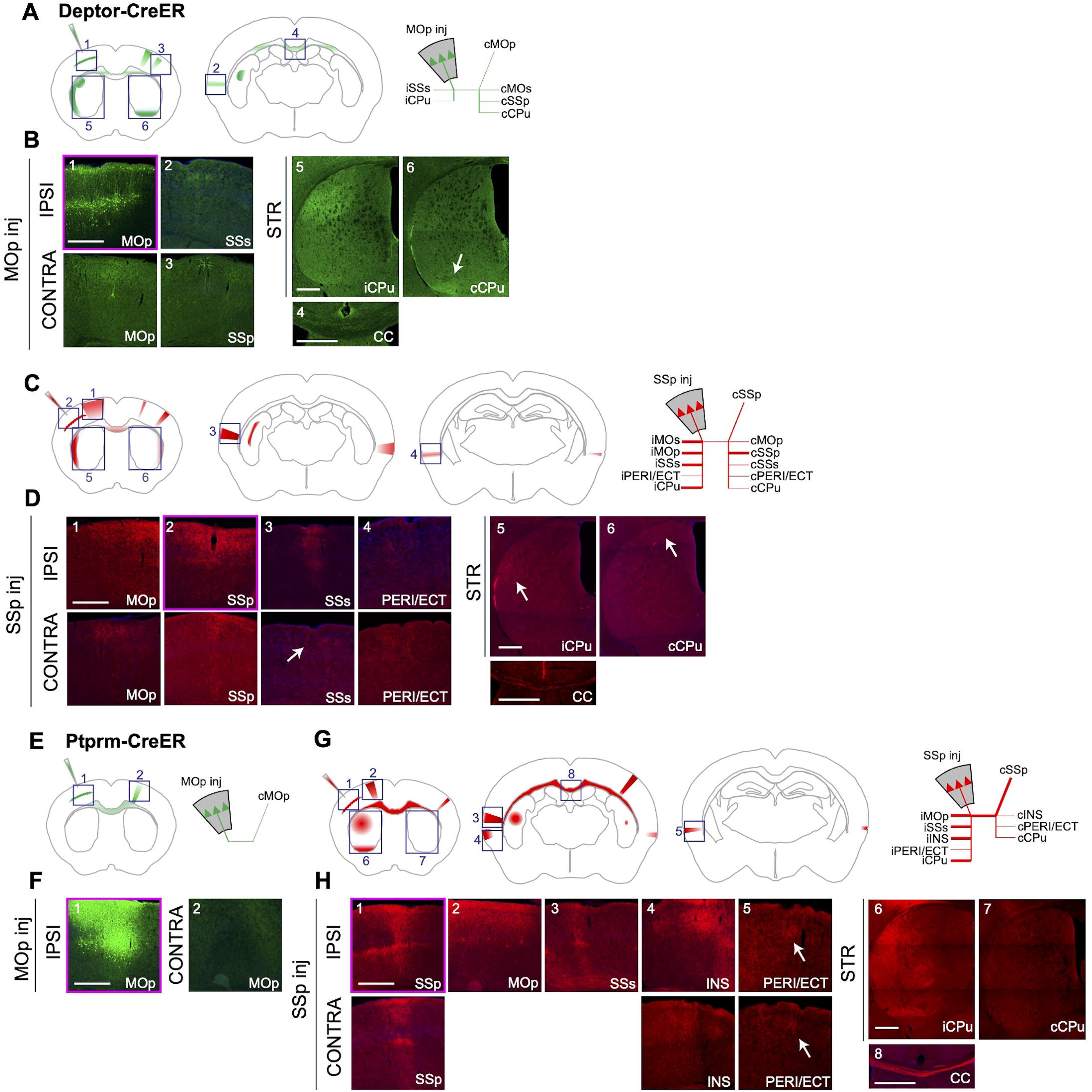
*Deptor-CreER* and *Ptprm-CreER* label distinct lower-layer IT neuron populations with contrasting long-range projection patterns. **(A,B)** EGFP-labeled anterograde projections from MOp of *Deptor-CreER* mice. **(A)** Schematic illustration (left) of major *Deptor*^+^ axonal projections from MOp. Stick diagram (right) of ipsilateral and contralateral *Deptor*^+^ axonal projections from primary motor cortex. **(B)** High-magnification confocal images of the boxed regions in (A) demonstrating EGFP-positive anterograde projections from MOp (magenta outline) in *Deptor-CreER* mice. Sparse cortico-cortical projections are observed in ipsilateral SSs and contralateral homotopic MOp, as well as heterotopic SSp. Bilateral corticostriatal projections were observed from motor cortex, concentrated in dorsolateral striatum ipsilaterally and ventral striatum contralaterally. *Deptor*^+^ callosal axons are much sparser than for upper-layer *Wfs1*^+^ IT neurons (compare to Figure 3B). **(C,D)** tdTomato-labeled anterograde projections from SSp of *Deptor-CreER* mice. **(C)** Schematic illustration (left) of major *Deptor*^+^ axonal projections from SSp. Stick diagram (right) of major (thick lines) and minor (thin lines) axonal projections from SSp in *Deptor-CreER* mice. **(D)** High-magnification confocal images of the boxed regions in (C) demonstrating sparse tdToamto^+^ cortico-cortical projections from SSp (magenta outline) to ipsilateral MOp and SSs, as well as contralateral homotopic SSp and heterotopic MOp. Very scant projections are observed in contralateral SSs and bilateral PERI/ECT. Sparse bilateral *Deptor*^+^ corticostriatal projections are observed from SSp. **(E,F)** EGFP-labeled anterograde projections from MOp of *Ptprm-CreER* mice. **(E)** Schematic illustration (left) and stick diagram (right), and **(F)** high-magnification confocal image of scant homotopic projections of *Ptprm*^+^ neurons from MOp (magenta outline) in *Ptprm-CreER* mice. **(G,H)** tdTomato-labeled anterograde projections from SSp of *Ptprm-CreER* mice. **(G)** Schematic illustration (left) of major *Ptprm*^+^ axonal projections from SSp (magenta outline) in *Ptprm-CreER* mice. Stick diagram (right) of major (thick lines) and minor (thin lines) axonal projections from SSp in *Ptprm-CreER* mice. **(H)** High-magnification confocal images of the boxed regions in (G) demonstrating ipsilaterally biased projections to MOp, SSs, INS, and PERI/ECT. Contralateral projections are observed in homotopic SSp, with sparser labeling observed in contralateral INS and PERI/ECT. Corticostriatal projections also demonstrate an ipsilateral projection bias with scant projections to contralateral lateral and ventral striatum. All anterograde projections were confirmed through observation across N=3 mice per line. IPSI = ipsilateral, CONTRA = contralateral, i = ipsilateral, c = contralateral, inj = injection, MOp = primary motor cortex, MOs = secondary motor cortex, SSp = primary somatosensory cortex, SSs = secondary somatosensory cortex, INS = insular cortex, PERI/ECT = perirhinal/ectorhinal cortex, CPu = caudate/putamen, CC = corpus callosum. Scalebars = 500 µm.

*Deptor*^+^ IT neurons in SSp demonstrated less dense callosal projections than *Wfs1*^+^ upper-layer IT neurons, but denser callosal projections than *Deptor*^+^ lower-layer IT neurons from MOp (compare Figure 3B,D with 4B,D). *Deptor*^+^ IT neurons in SSp sent dense projections to ipsilateral MOp and SSs. In contralateral cortex, they primarily targeted homotopic contralateral SSp with low density heterotopic projections to contralateral MOp and SSs. Very sparse projections were observed in bilateral PERI/ECT. Across laminae in all targeted cortical areas, *Deptor*^+^ axons were most dense in layer 2/3. Corticostriatal projections of *Deptor*^+^ IT neurons from SSp differed from those in MOp by their symmetry, demonstrating dense ipsilateral projections to dorsolateral and caudal striatum, with more diffuse, non-localized contralateral corticostriatal projections (Figure 4D).

### Ptprm-CreER

*Ptprm* is a type IIB receptor protein tyrosine phosphatase. Like Igfn1, it contains immunoglobulin-like and fibronectin type III domains (Hay et al., 2023). Ptprm is a transmembrane protein involved in cell adhesion (Hay et al., 2023). When evaluating only excitatory projection neurons of the isocortex, *Ptprm* is expressed in a narrow band of layer 5 IT neurons (Figure 1C’, Figure 5D, Supplementary Figures 1 and 2). However, *Ptprm* expression is not exclusive to excitatory neurons, with expression in cortical interneurons and endothelial cells (Yao et al., 2023).

**Figure 5:**
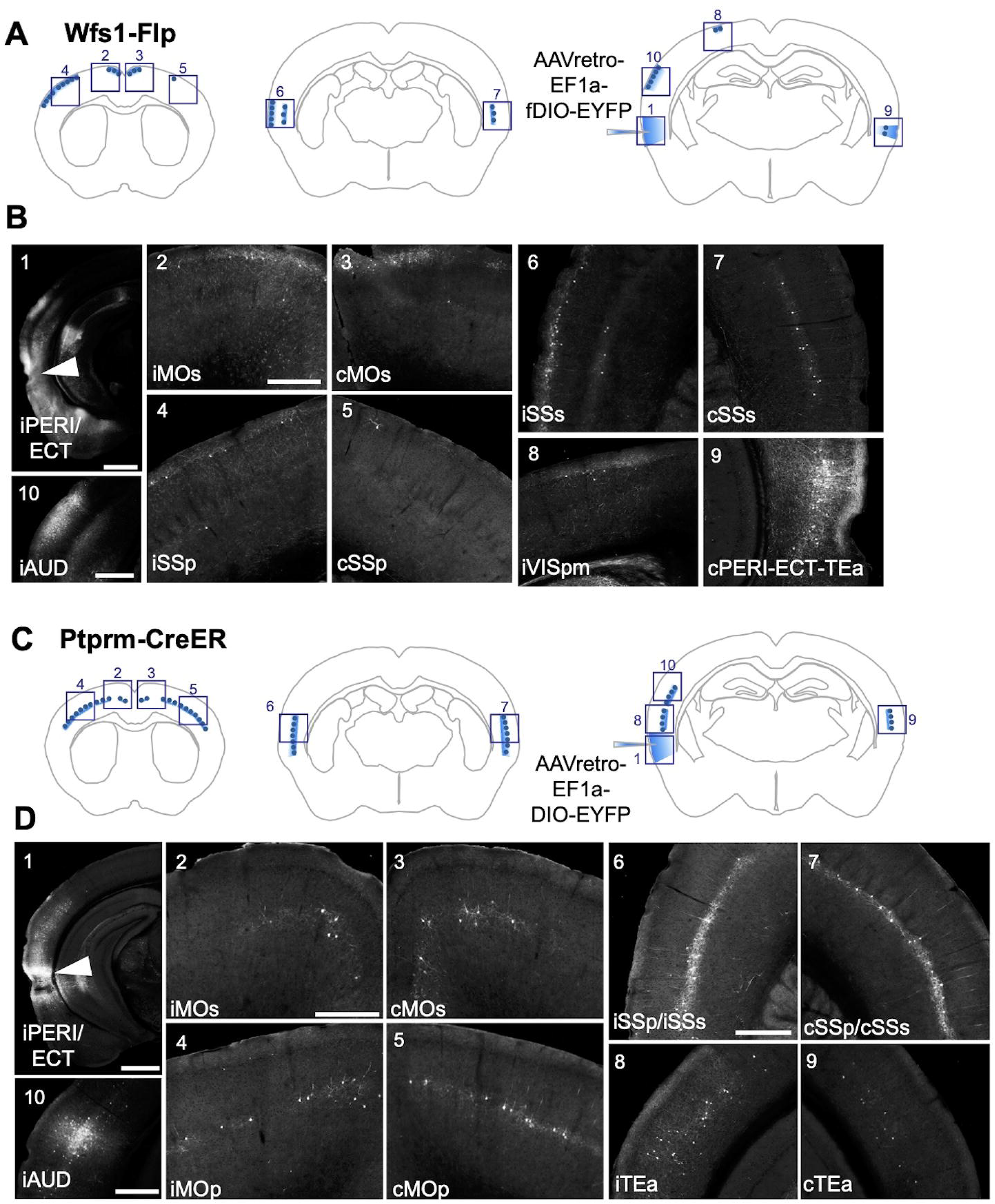
Comparable areal input to perirhinal/ectorhinal cortex from upper-layer *Wfs1*^+^ and lower-layer *Ptprm*^+^ IT neurons. **(A)** Schematic representation of Flp-dependent EYFP packaged in an AAVretro capsid stereotaxically injected into PERI/ECT and densities of retrogradely-labeled *Wfs1*^+^ IT neurons in *Wfs1-Flp* mice. **(B)** Representative image of the PERI/ECT injection site (arrowhead), as well as EYFP^+^ retrogradely-labeled cell bodies, which were primarily observed in bilateral MOs and SSs, as well as ipsilateral AUD and contralateral PERI/ECT/TEa. Very sparse retrograde labeling was noted in bilateral MOp, SSp and ipsilateral VISpm. Note the laminar differences in retrograde cell body labeling in ipsilateral versus contralateral SSs, where *Wfs1*^+^ neurons in L2/3 and L5 project to ipsilateral PERI/ECT but only *Wfs1*^+^ neurons in L5 project to contralateral PERI/ECT. **(C)** Schematic representation of Cre-dependent EYFP packaged in an AAVretro capsid stereotaxically injected into PERI/ECT and densities of retrogradely-labeled *Ptprm*^+^ IT neurons in *Ptprm-CreER* mice. **(D)** Representative image of the PERI/ECT injection site (arrowhead), as well as EYFP^+^ retrogradely-labeled cell bodies, which were primarily observed in bilateral SSs with lower density labeling in bilateral MOs, MOp, SSp, and TEa. Dense labeling was also observed in ipsilateral auditory cortex. The primary areal differences in PERI/ECT-projecting upper-layer *Wfs1*^+^ versus lower-layer *Ptprm*^+^ IT neurons is from ipsilateral *Wfs1*^+^ IT neurons in visual cortex and bilateral *Ptprm*^+^ IT neurons in TEa. All retrograde projections were confirmed through observation across N=3 mice per line. i = ipsilateral, c = contralateral, MOp = primary motor cortex, MOs = secondary motor cortex, SSp = primary somatosensory cortex, SSs = secondary somatosensory cortex, INS = insular cortex, PERI/ECT = perirhinal/ectorhinal cortex, TEa = temporal association areas, AUD = auditory cortex, VISpm = posteromedial visual area. Scalebars for panels B1 and D1 (injection sites) = 1000 µm, all other scalebars = 500 µm. Scalebars in panel B2 and D2 apply to panels 2-9.

To explore the projection patterns of *Ptprm*^+^ IT neurons, we generated a transgenic knock-in mouse line that introduces a T2A-IRES-CreER^T2^ between the final exon of *Ptprm*, exon 31, and its 3’ UTR. Crossing *Ptprm-CreER* with the Cre-dependent *Ai14* reporter line (Madisen et al., 2010), we observed tdTomato reporter expression in excitatory layer 5 pyramidal neurons, as well as interneurons and blood vessels scattered throughout the cortex (Figure 1C-D). Subcortically, *Ptprm*-driven reporter expression was also apparent in the olfactory bulb, and several thalamic, hypothalamic, and amygdalar nuclei (Supplementary Figure 4E,F).

Anterograde AAV2/9 tracing with Cre-dependent reporters from MOp and SSp was then used to investigate projection patterns of *Ptprm*^+^ layer 5 IT neurons (Figure 4E-H). Because cortical interneurons and endothelial cells are locally restricted, any long-range *Ptprm*^+^ axonal projections can be inferred to arise from layer 5 IT neurons. Projection patterns from *Ptprm^+^* IT neurons varied by cortical area. Nearly no anterograde projections were observed from MOp, with only sparse callosal projections to homotopic MOp and no other notable cortco-cortical connections.

In SSp, on the other hand, *Ptprm*^+^ IT neurons exhibited dense callosal projections to layers 1, 2/3, and 5 of homotopic contralateral SSp, as well as ipsilateral projections to layers 1, 2/3, and 5 of MOp, SSs, and INS, with less prominent projections to ipsilateral PERI/ECT. Very sparse projections were observed to contralateral INS and PERI/ECT. Corticostriatal projections of *Ptprm*^+^ IT neurons from SSp differed from those of *Deptor*^+^ IT neurons. *Ptprm*^+^ IT neurons in SSp exhibited dense ipsilateral projections to dorsolateral striatum, ventral striatum, and caudal striatum, but comparatively few projections to contralateral striatum. Scant reporter positive striae were also noted passing through the striatum and internal capsule suggesting that a minor subset of ET neurons also express *Ptprm*, consistent with single-cell expression data (Supplementary Figure 1C).

### IT neuron projections to PERI/ECT

Both upper-layer *Wfs1*^+^ and lower-layer *Ptprm*^+^ IT neurons in sensorimotor cortex exhibited notable projections to PERI/ECT, which is a point of broad cortical convergence and an entry point to the hippocampal formation, mPFC, and ENTl as part of the posterolateral temporal subnetwork (Zingg et al., 2014). Although several studies focus on cortico-cortical projections to PERI/ECT from rodent motor cortex (Munoz-Castaneda et al., 2021; Ueta et al., 2019), extensive bidirectional tracing illustrates that PERI/ECT is densely interconnected with most isocortical areas. PERI/ECT receives input from neurons in layers 2 and 5a across cortical areas (Zingg et al., 2014), corresponding to the laminar distributions of *Wfs1*^+^ and *Ptprm*^+^ IT neurons, respectively. Moreover, along the anterior-posterior axis, the “middle” PERI/ECT is more interconnected with sensorimotor subnetworks, as well as visual and auditory subnetworks (Zingg et al., 2014). We therefore performed retrograde viral tracing from middle PERI/ECT in both *Wfs1-Flp* and *Ptprm-CreER* lines to catalog subtype-specific differences in cortico-cortical input to PERI/ECT (Figure 5).

A retrograde AAV carrying a Flp- or Cre-dependent EYFP reporter was unilaterally stereotaxically injected into coordinates centered around middle PERI/ECT in adult *Wfs1-Flp* or *Ptprm-CreER* mice, respectively. Cre-dependent EYFP expression in *Ptprm-CreER* mice was subsequently induced with tamoxifen, and STPT was used in both lines to identify EYFP-labeled PERI/ECT-projecting IT neuron somata across cortical areas.

Retrogradely labeled *Wfs1*^+^ neurons were identified in ipsilateral and contralateral isocortex, most prominently in ipsilateral auditory cortex (AUD), as well as contralateral PERI/ECT and temporal association area (TEa), suggesting dense interconnectivity between bilateral PERI/ECT/TEa. We also observed retrogradely labeled *Wfs1*^+^ neurons in bilateral MOs and SSs, and very sparse retrograde labeling from bilateral MOp, bilateral SSp, and ipsilateral posteromedial visual area (VISpm). Of note, there was a clear asymmetry of laminar labeling in SSs with ipsilateral *Wfs1*^+^ neurons residing mostly in layer 2/3 and contralateral Wfs1^+^ neurons residing entirely in layer 5 (Figure 5B).

Retrogradely labeled *Ptprm*^+^ neurons also exhibited the highest densities in ipsilateral AUD and bilateral SSs. *Ptprm*^+^ IT neurons were also observed at slightly lower density in bilateral MOs, MOp, SSp, and TEa, though contralateral TEa was sparser than ipsilateral TEa. The main areal distinctions between the IT lines is that primary sensorimotor cortices contain a greater density of lower-layer *Ptprm*^+^ IT neurons than upper-layer *Wfs1*^+^ IT neurons that project to PERI/ECT, consistent with prior retrograde tracing data that shows a greater density of layer 5a than layer 2 neurons projecting to TEa (and associated PERI/ECT) from the more medial limb-associated regions of SSp (Zingg et al., 2014). We also observe that PERI/ECT receives input from contralateral *Wfs1*^+^ TEa neurons versus ipsilateral *Ptprm*^+^ TEa neurons. Finally, bilateral MOs, SSs, and ipsilateral auditory cortex have similar densities of upper-layer *Wfs1*^+^ and lower-layer *Ptprm*^+^ PERI/ECT projecting IT subtypes.

### In vivo functional responses of contrasting IT subtypes in a reach-withdraw-drink task

To investigate the differential *in vivo* electrophysiology of upper-versus lower-layer IT neurons during motivated behavior, we imaged calcium-dependent activity in upper-layer *Wfs1*^+^ or lower-layer *Deptor*^+^ IT neurons in the forelimb subregion of SSp (SSp-ul) of head-restrained mice. To perform these experiments, *Wfs1-Flp* mice received an SSp injection of Flp-dependent GCaMP8fv, while *Deptor-CreER* mice were crossed with GCaMP8m mice to generate *Deptor-iCreER;GCaMP8m* mice that were subsequently induced with tamoxifen to activate *GCaMP8m* expression. Mice performed a single-handed RWD task in the dark (Figure 6A,B). In this previously established paradigm (Galinanes et al., 2018; Li et al., 2026), mice use chemosensory and vibrissae cues to locate a waterspout presented at one of three positions spanning the peri-snout space (P1: ipsilateral to the reaching limb; P2: center; P3: contralateral), then reach, grasp the water droplet, and withdraw the hand to the mouth for consumption. Trained mice adopt distinct forelimb trajectories depending on waterspout location while maintaining accurate grasp endpoints across positions, consistent with prior characterization of this behavior (Li et al., 2026).

**Figure 6:**
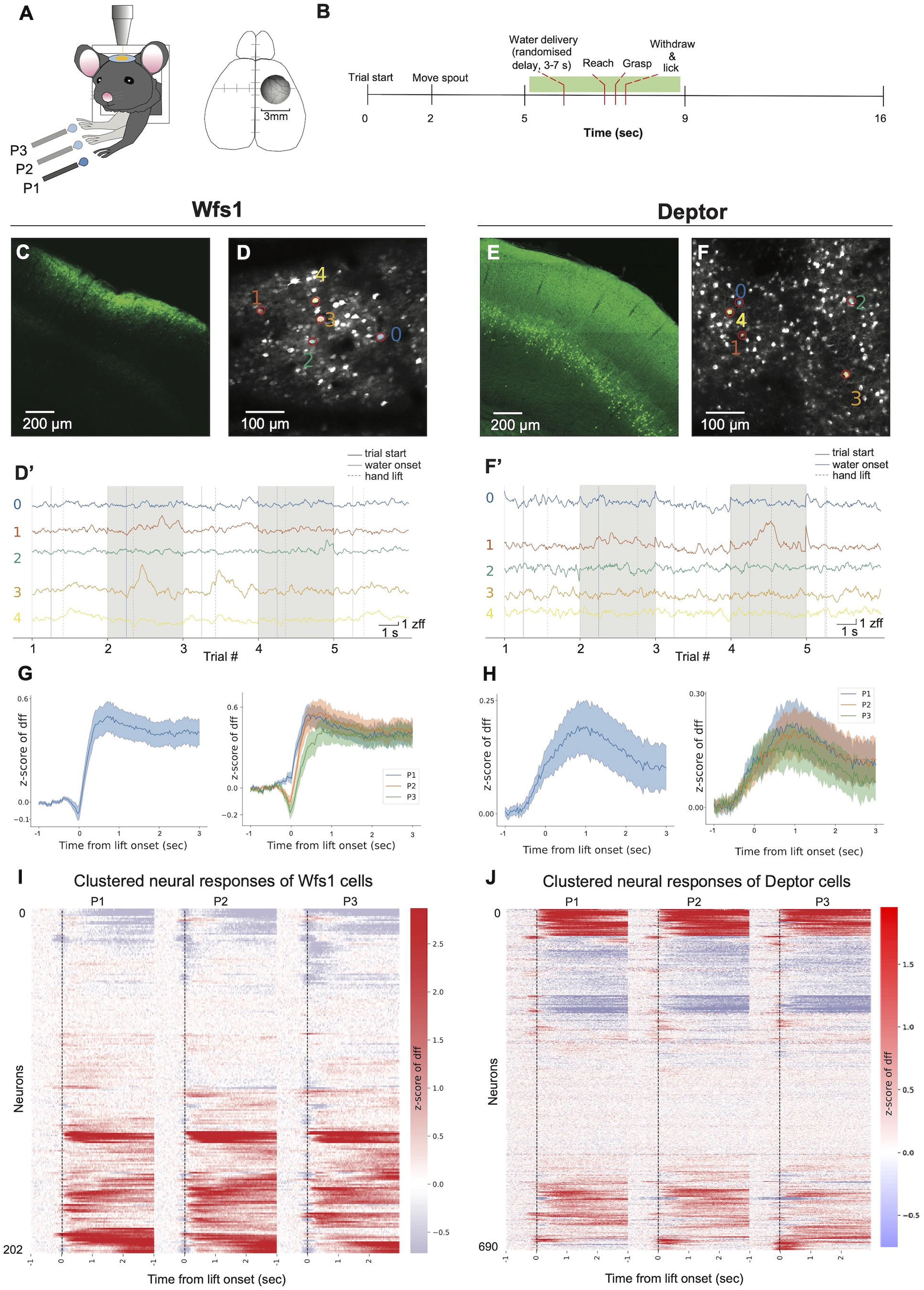
*In vivo* calcium imaging reveals temporally distinct responses of upper-layer *Wfs1*+ IT neurons compared to lower-layer *Deptor*+ IT neurons in a reach-withdraw-drink task. **(A)** Schematic of a head-restrained mouse (left) performing a reach-withdraw-drink task and reaching with the left forelimb toward a waterspout positioned at one of three locations (ipsilateral P1, central P2, contralateral P3). Schematic of the 3mm two-photon cranial imaging window (right) centered over the right SSp-ul. **(B)** Timeline for one trial in the reach-withdraw-drink task. **(C)** Confocal image confirming cell-type-specific GCaMP expression in SSp-ul in a *Wfs1-Flp* mouse injected with a Flp-dependent GCaMP8f virus. **(D)** Maximum projection image of a representative FOV from a recording session of a *Wfs1-Flp* mouse injected with Flp-dependent GCaMP8fv. **(D’)** Individual ΔF/F₀ traces corresponding to the five example ROIs highlighted in (D) during five random single trials. **(E)** Confocal image confirming cell-type-specific GCaMP expression in SSp-ul in a *Deptor-iCreER;GCaMP8m* reporter mouse. **(F)** Maximum projection image of a representative FOV from a recording session in a *Deptor;GCaMP8m* mouse. **(F’)** Individual ΔF/F₀ traces corresponding to the five example ROIs highlighted in (F) during five random single trials. **(G)** Average calcium activity of SSp-ul neurons in Wfs1-Flp mice injected with Flp-dependent GCaMP8fv. Average activity is aligned to waterspout contact. Left: activity pooled across all spout locations. Right: activity separated by spout location (P1, P2, P3). Data are collected from 202 *Wfs1*^+^ neurons from N=4 mice. Shading indicates mean ± standard error. **(H)** Average calcium activity of SSp-ul neurons in *Deptor;GCaMP8m* mice aligned to waterspout contact. Left: activity pooled across all spout locations. Right: activity separated by spout location (P1, P2, P3). Data are collected from 690 *Deptor*^+^ neurons from N=4 mice. Shading indicates mean ± standard error. **(I)** Hierarchically clustered heatmap of activity from individual *Wfs1*^+^ GCaMP8fv-expressing neurons averaged across trials and aligned to forelimb lift onset, organized by spout location (from left to right: P1, P2, P3). Color scale indicates ΔF/F₀. **(J)** Hierarchically clustered heatmap of individual *Deptor;GCaMP8m* neuron activity averaged across trials and aligned to forelimb lift onset, organized by spout location (from left to right: P1, P2, P3). Color scale indicates ΔF/F₀.

Using two-photon calcium imaging through a cranial window over SSp-ul (Figure 6A, C-F’), we recorded GCaMP8fv fluorescence in *Wfs1*^+^ neurons and GCaMP8m fluorescence in *Deptor*^+^ neurons. Cell type-specific expression was confirmed by post hoc histological validation of the respective recording sites (Figure 6C,E). Representative field of view (FOV) images from both lines showed robust labeling of distinct IT populations with their expected laminar distributions, and single-cell calcium traces exhibited clear fluorescence transients across individual trials (Figure 6D’,F’), validating that both IT subtypes could be reliably monitored *in vivo*.

Population-level activity averaged across all recorded SSp-ul neurons revealed markedly different response profiles between the two IT neuron subtypes (Figure 6G,H). *Wfs1*^+^ neurons displayed a pronounced suppression at movement onset, returned toward baseline near the time of spout touch, and peaked around hand lick (Figure 6G), demonstrating that *Wfs1*^+^ neuronal activity may be precisely gated by discrete behavioral events. Moreover, *Wfs1*^+^ neurons showed graded, location-dependent responses that distinguished ipsilateral from central reach directions (Figure 6G). In contrast, *Deptor*^+^ neurons exhibited a slow, sustained ramp in activity following waterspout contact that lacked a discrete peak and showed no spout location selectivity (Figure 6H), implying that *Deptor*^+^ IT neurons are tonically recruited throughout the reach without encoding the directional demands of the movement.

At the single-neuron level, hierarchical clustering of trial-averaged activity revealed further distinctions between the two populations (Figure 6I,J). *Wfs1*^+^ neurons displayed temporally precise and heterogeneous response profiles, including clusters exhibiting brief peri-event suppression (Figure 6I). *Deptor*^+^ neurons, in contrast, were dominated by broad, persistent increases or decreases in activity spanning the duration of the behavior, implying a role in continuous sensorimotor monitoring of limb movement (Figure 6J). Taken together, these results establish that upper-layer *Wfs1*^+^ and lower-layer *Deptor*^+^ IT subtypes in SSp-ul are differentially recruited during skilled forelimb behaviors, with *Wfs1*^+^ IT neurons encoding discrete, directionally selective sensorimotor events and *Deptor*^+^ IT neurons providing sustained, trajectory-level signals.

## DISCUSSION

An important milestone in cortical evolution was the expansion of indirect over direct neurogenesis, whereby increasingly large subsets of excitatory neurons in dorsal pallium were generated from intermediate progenitors rather than directly from radial glia cells (Cardenas & Borrell, 2020; Deryckere et al., 2025). The increased proportion of indirect neurogenesis in higher-order mammals served to amplify and diversify intratelencephalic over extratelencephalic excitatory projection neurons, which is thought to be a critical factor underlying the increased cognitive capacity of primates and humans (Cardenas & Borrell, 2020; Greig et al., 2013; Huilgol et al., 2025; Tosches & Laurent, 2019). Single-cell profiling techniques have expanded our appreciation for the diversity of IT subpopulations at the transcriptomic (Fang et al., 2022), anatomic (Gao et al., 2022; Peng et al., 2021), and electrophysiologic levels (Berg et al., 2021). However, distinct IT subtypes remain poorly defined, in part because of a lack of tools that link marker expression with axonal projection patterns in different IT subsets across laminae. Cortical profiling studies rely on a small battery of classic IT markers, including *Satb2*, *Cux1*, *Cux2*, *Lhx2*, and *Plxnd1* (Matho et al., 2021; Munoz-Castaneda et al., 2021; Peng et al., 2021), which, although generally restricted to the IT class, have broad translaminar expression and do not isolate IT subtypes with unique cortico-cortical and corticostriatal projections. We therefore aimed to expand the repertoire of IT-specific tools by introducing six new knock-in Flp or CreER mouse driver lines and using them to characterize the long-range projection patterns and electrophysiological activity of distinct molecularly defined IT subpopulations.

These mouse lines facilitate easy genetic access to IT subsets whose projection motifs could not previously be linked through a unified molecular and hodological definition. For example, (Yamashita et al., 2018) describe layer 2/3 neurons in SSp that project to ipsilateral MOp/MOs, SSs, TEa/PERI/ECT, and dorsolateral striatum, consistent with the projection patterns of *Wfs1*^+^ neurons in SSp in our study. Similarly, using mixed retrograde tracing techniques, Muñoz-Castaneda, et al. (2021) also identify IT neurons in layer 2 of MOp that project to bilateral TEa/PERI/ECT, mirroring our observations for *Wfs1*^+^ IT neurons. Likewise, PERI-projecting neurons from the *Wfs1*-rich layer 2 of rat MOs have been described (Ueta et al., 2013; Ueta et al., 2019), consistent with the *Wfs1*^+^ cell bodies in MOs we observed in mice through retrograde tracing from PERI/ECT. *Rasgrf2* labels upper-layer IT neurons with similar projection motifs (Liu et al., 2024), on par with the layer 2/3 population labeled by *Plxnd1* (Matho et al., 2021), both of which span layer 2/3 more broadly than *Wfs1*, which remains largely restricted to superficial-most layer 2.

*Igfn1*^+^ IT neurons, whose cell bodies reside deeper in layer 2/3 than *Wfs1*^+^ neurons, were surprising in the sparseness of their long-range projections. Prior literature provides evidence for locally-projecting upper-layer IT neurons without long-range projections. For example, certain layer 2/3 non-callosal neurons make only intrahemispheric projections from SSp to MOp (Yamashita et al., 2018). Likewise, a subcluster of non-callosal, non-corticostriatal IT neurons has been described in lower layer 2/3 and layer 4 of MOp that project only to ipsilateral MOs or SSp (Munoz-Castaneda et al., 2021). In primary visual cortex (V1), *Igfn1* is expressed in an IT subpopulation whose transcriptomic signature only emerges with visual input (Cheng et al., 2022), suggesting that *Igfn1*^+^ neurons may alter features of their identity in a sensory-dependent manner. Future work investigating whether *Igfn1*^+^ IT neurons rely on sensory stimuli in other primary cortical sensory areas, and to what extent sensory-dependent gene expression alters the properties of *Igfn1*^+^ neurons, may reveal fundamental principles of how IT neurons adapt to emerging sensory input during critical periods of plasticity in early cortical development.

Existing literature is less explicit about specific subtypes of lower-layer IT neurons, since most compare layer 5 IT neurons with ET neurons (Baker et al., 2018; Kalmbach et al., 2021). However, a few specifically detail the projection patterns of molecularly defined lower-layer IT neurons. For example, Im et al., (2022) describe a population of *Tlx3^+^* IT neurons in layer 5a of mouse MOs that preferentially target layer 1 of ipsilateral AUD and PERI, extend axons into the corpus callosum, and target both dorsal and ventral striatum. In SSp, *Tlx3*^+^ IT neurons also illustrate broad bilateral cortico-cortical and corticostriatal projections (Liu et al., 2024). Kim et al., (2015) contrast *Tlx3-Cre* and *Efr3a-Cre* lines to distinguish two subpopulations of layer 5 IT neurons in mouse V1, observing that *Tlx3*^+^ IT neurons project to ipsilateral secondary visual cortex (V2), contralateral homotopic V1, and ipsilateral striatum, while *Efr3a*^+^ neurons only project locally to ipsilateral adjacent cortical areas. The lower-layer *Ptprm*^+^ subpopulation we describe in SSp shares some characteristics with the *Tlx3*^+^ population described in MOs and V1, including projections to ipsilateral PERI, callosal projections, and ipsilateral corticostriatal projections that span both dorsal and ventral striatum, though *Ptprm*^+^ IT neurons appear to have an ipsilateral projection preference compared to the bilateral projections of *Tlx3*^+^ IT neurons (Liu et al., 2024). Thus, our lines enhance the breadth of tools available to untangle the unique properties of more narrowly defined lower-layer IT neurons.

We observed projections from bilateral sensorimotor cortex to PERI/ECT in multiple IT lines, though they were most prominent for upper-layer *Wfs1*^+^ IT neurons and lower-layer *Ptprm*^+^ IT neurons. Perirhinal cortex, which is part of the posterolateral temporal subnetwork (Zingg et al., 2014), is thought to serve as an “associative hub” for multimodal cortico-cortical sensory input, binding relevant sensory features for memory encoding by the hippcocampal-parahippocampal network (Holmes et al., 2026; Zhang & Zhang, 2026). It is therefore interesting to note specific distinctions in IT projections from primary and secondary cortical areas that vary by upper- or lower-layer IT subtype. While secondary sensorimotor areas exhibit similar densities of PERI/ECT-projecting upper-layer *Wfs1*^+^ and lower-layer *Ptprm*^+^ IT neurons, SSp contains more lower-layer *Ptprm*^+^ than upper-layer *Wfs1*^+^ IT neurons that project to PERI/ECT, in line with prior observations of less dense layer 2 input from medial limb-associated SSp (Zingg et al., 2014). Additionally, PERI/ECT receives input from contralateral *Wfs1*^+^ neurons in TEa, in contrast with ipsilateral-predominant input from *Ptprm*^+^ neurons in TEa.

In addition to cortico-cortical connections, IT neurons also project to the striatum. Generally, upper-layer IT neurons favor ipsilateral corticostriatal projections while lower-layer IT neurons favor bilateral corticostriatal projections (Peng et al., 2021; Shepherd, 2013; Wang et al., 2026), motifs that are broadly conserved across most of our lines. However, despite the use of consistent stereotaxic injection coordinates for anterograde tracing, we observed marked variability in the laterality, symmetry, and regional targeting of corticostriatal projections across IT lines. Recent striatal projectome atlases demonstrated that corticostriatal projections from sensorimotor cortex were largely organized into a somatotopic map in the lateral caudoputamen (Hintiryan et al., 2016; Wang et al., 2026). Corticostriatal projections from our IT lines reflect the known anatomy of projections from forelimb sensorimotor cortex to the forelimb region of the ipsilateral lateral caudoputamen (Hintiryan et al., 2016; Wang et al., 2026). Subtype-specific differences arise, however, in their contralateral symmetry and projections to other striatal subdivisions. For example, Wang et al., (2026) suggest that layer 5 IT neurons exhibit the strongest convergence on striatal subdivisions. However, we observe divergent lower-layer corticostriatal projection patterns: *Deptor*^+^ IT neurons in SSp project to a diffuse dorsoventral region of the contralateral striatum with a prominent ventral striatal component, while *Ptprm*^+^ IT neurons in SSp make few contralateral striatal projections.

A major strength of our study is that we expand the breadth of mouse lines available for future investigations of cortical organization and IT neuron physiology. Moreover, we strategically selected Flp-dependent expression for upper-layer IT lines and CreER-dependent expression for lower-layer IT lines to enable future intersectional strategies that allow upper- and lower-layer IT populations to be compared within the same individual animal. Intersecting pairs of IT lines may help dissect even narrower and more specific IT subtypes through combinatorial labeling. Nevertheless, there remain limitations in using these lines to thoroughly catalog IT subtypes. First, the IT markers we selected vary in their breadth and the degree to which they overlap with each other (Supplementary Figure 2). Thus, these markers do not necessarily reflect mutually exclusive IT subpopulations. Moreover, these markers are not exhaustive of all potential IT subtypes, though a definitive catalog would depend on what multi-modal neuronal properties are used to define bona fide IT subtypes. For example, we did not develop a line to isolate rare layer 6 IT neurons, though *Sulf1* (Munoz-Castaneda et al., 2021; Extended Data Fig 9) may be a good contender, as is *Oprk1* which labels lower-layer claustrum-like cortical neurons (Fang et al., 2025). Finally, our methods evaluate each IT line at the population level, but single-cell profiling studies of cortical IT neurons have reported variable projection patterns even within transcriptomic subclasses (Berg et al., 2021; Peng et al., 2021; Ueta et al., 2019; Yamashita et al., 2018), suggesting there may be even finer hodological distinctions to be made for individual IT neurons within our labeled subpopulations.

Our mouse lines enhance the richness of future experiments that dissect how IT neurons participate in cortical function and dysfunction. Looking forward, future use of these lines will help illuminate how IT neurons are incorporated into cortical microcircuits and modulate information flow. Finally, these markers are also expressed in human cortex (https://brain-map.org/our-research/cell-types-taxonomies/cell-types-database-rna-seq-data#human; https://human.brain-map.org/ish/search), and it remains to be seen whether they label homologous cell types with consistent projection patterns to what we describe in mice.

## MATERIALS AND METHODS

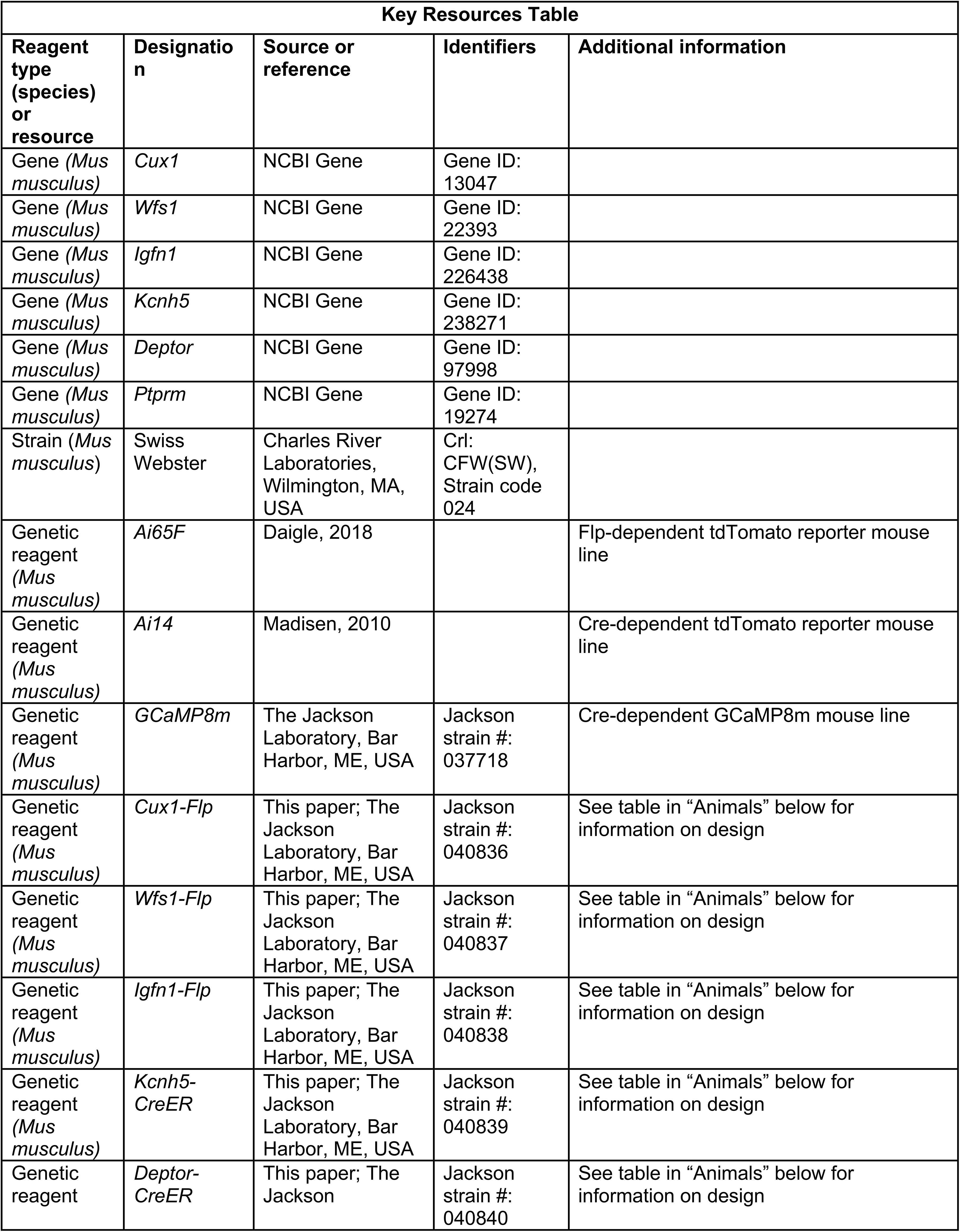
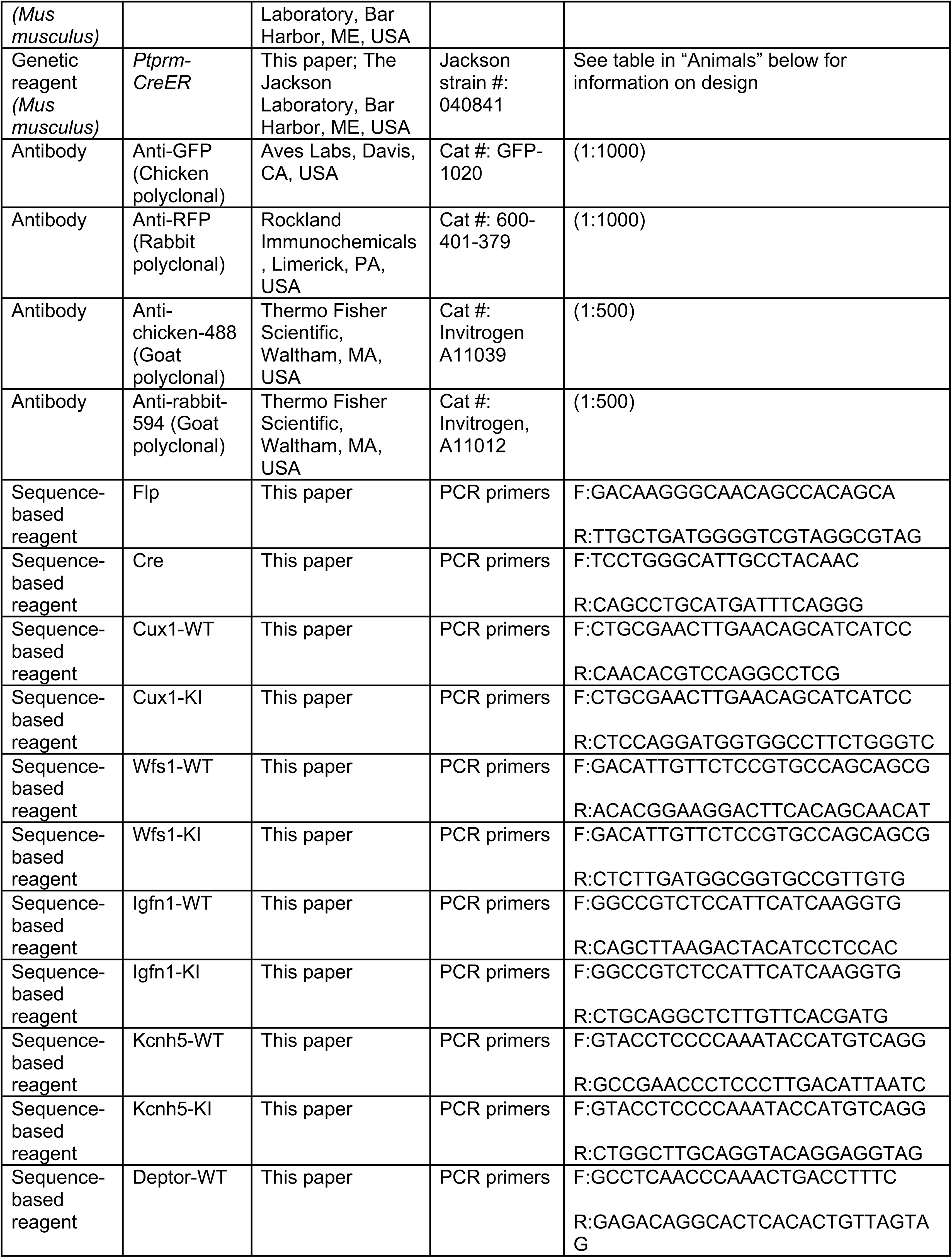
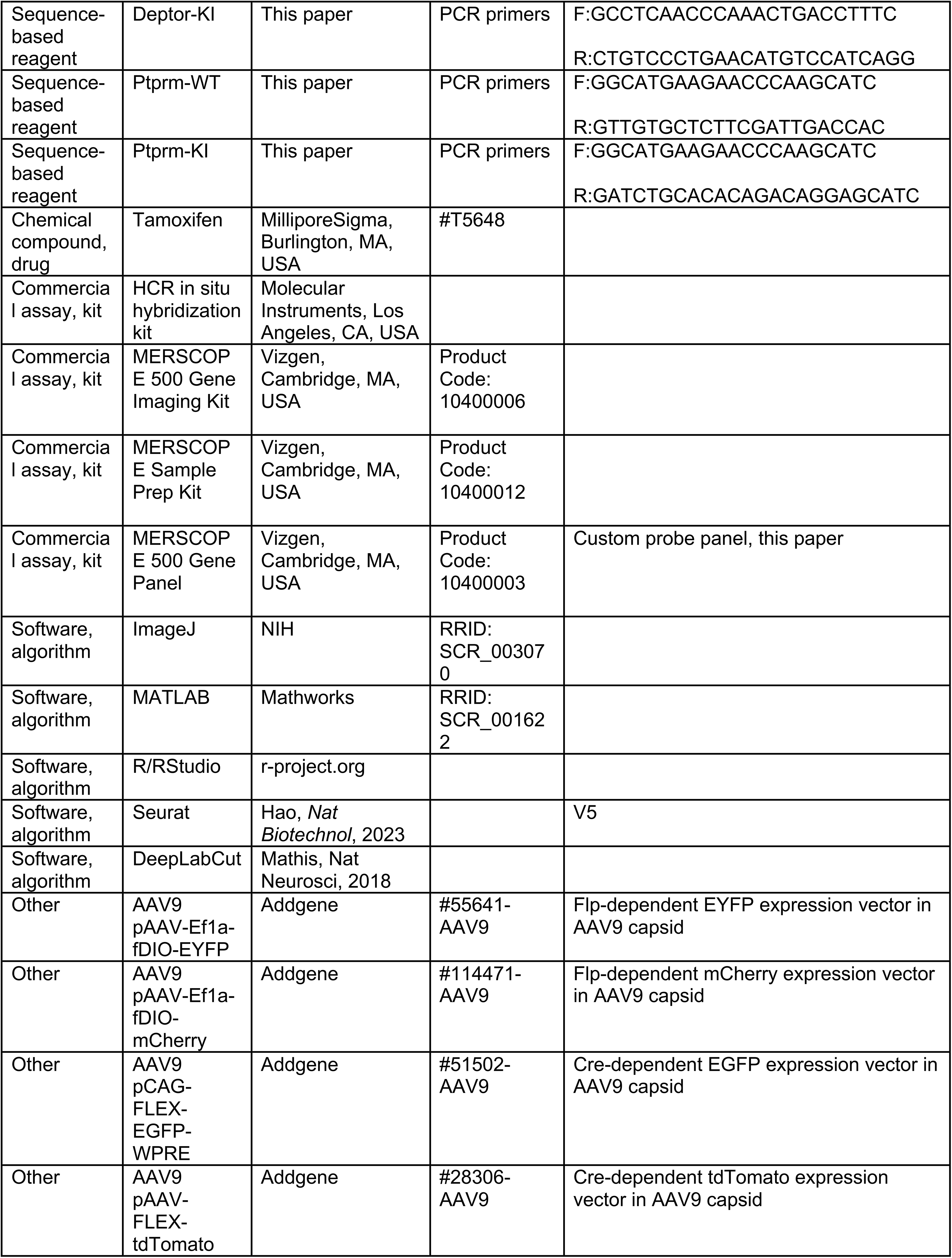
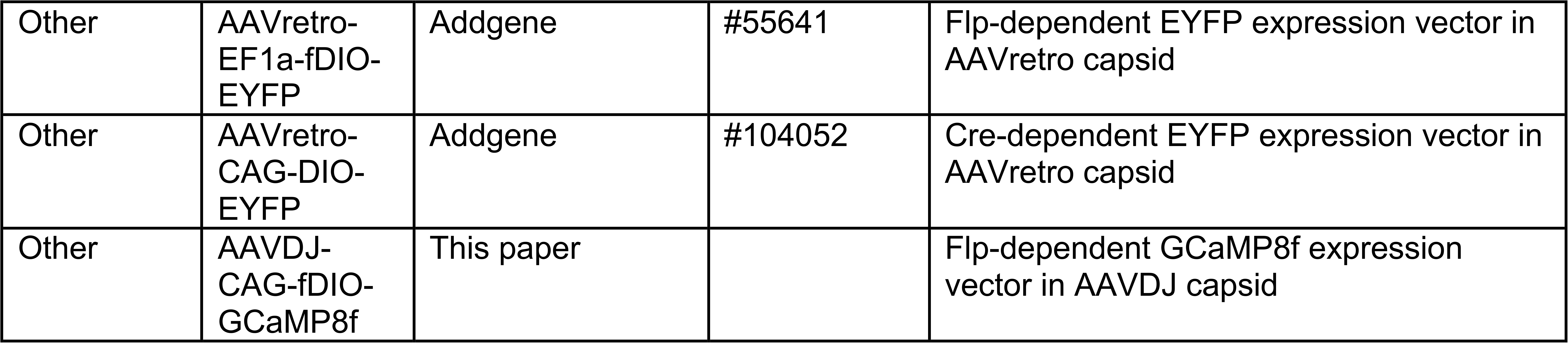

### Animals

All novel knock-in mouse lines reported in this manuscript were generated commercially by Biocytogen (Beijing, China) using their CRISPR/Cas9-based Extreme Genome Editing System. All mouse lines have been deposited for distribution at Jackson Laboratories (see table for stock numbers). Specific knock-in mouse line design features are as follows:

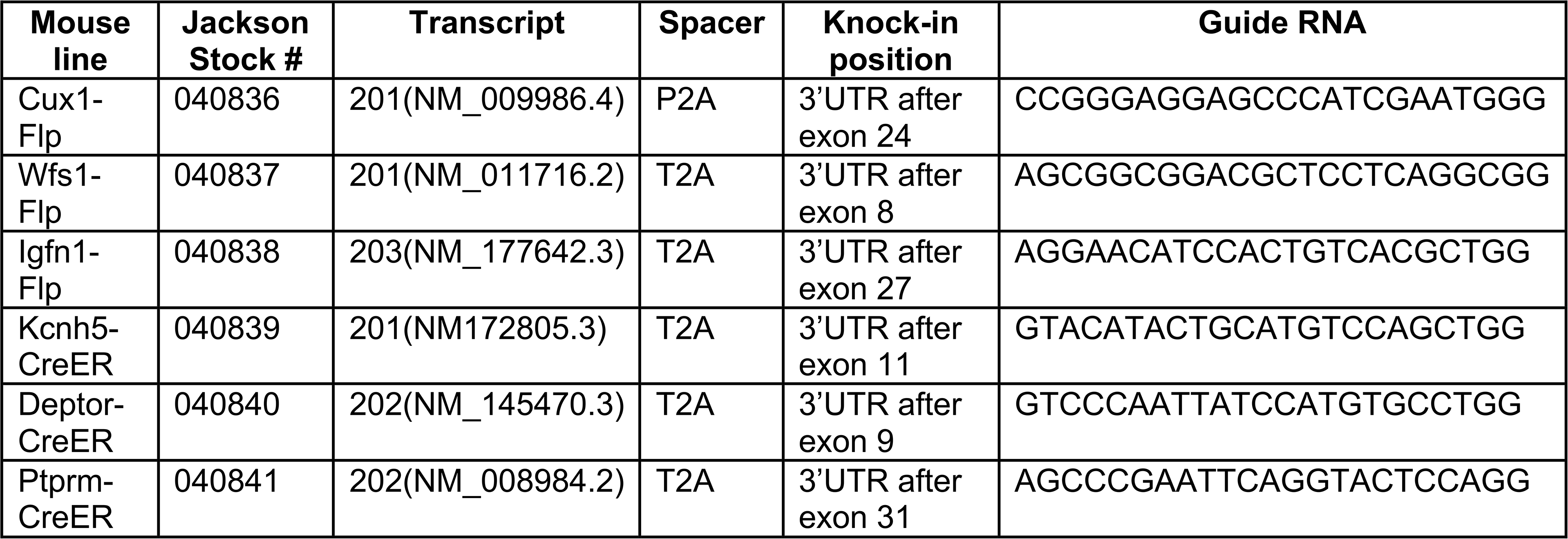

Additional lines used in these experiments include the Flp-dependent tdTomato Ai65F reporter line (Daigle et al., 2018), the Cre-dependent tdTomato Ai14 reporter line (Madisen et al., 2010), the Cre-dependent GCaMP8m line (The Jackson Laboratory, Bar Harbor, ME, USA), and wild type Swiss Webster (Charles River Laboratories, Wilmington, MA, USA) mice. All vertebrate animal work was performed in compliance with Duke IACUC guidelines. N ≥ 3 animals, both males and females, were included in every analysis.

### Brain tissue harvesting

Mice were anesthetized with isoflurane and transcardially perfused with 1X PBS, followed by 4% paraformaldehyde (PFA). Brains were harvested and post-fixed in 4% PFA overnight, then cryoprotected in 30% sucrose in 1X PBS until fully equilibrated. Tissues were subsequently embedded in OCT, flash-frozen in isopentane with dry ice, and then stored at −80 °C prior to cryostat sectioning.

### Serial two-photon microscopy and quantification

Perfused and fixed brains were shipped in 1X PBS to the Neuroimaging and Behavior Core Facility at Cold Spring Harbor Laboratory for imaging via STPT as previously described (Kim et al., 2017; Matho et al., 2021). Brains were embedded in oxidized agarose and kept in a sodium borohydride solution for three hours at room temperature for crosslinking. Whole brains were serially imaged in coronal planes at 50µm intervals. For two-photon excitation, a coherent chameleon-ultra laser tuned at 920 nm was used to excite tdTomato and EYFP. 14×18 (x by y) tiled serial two-photon images were collected for each brain at 1µm x 1µm resolution. Images were then stitched as a grid sequence. Each brain was then registered to the Allen reference Common Coordinate Framework version 3 (CCFv3) using custom-built software previously described (Matho et al., 2021). Automated cell body detection was performed using a convolutional network as previously described (Matho et al., 2021) or a custom-built macro in Fiji/ImageJ (for datasets with dense and high fluorophore expression and mixed cell size distribution). A table for center of mass (x,y,z) for all identified cells in the brain was prepared. Spatial information of detected cells across the whole brain was then used to map it onto the registered Allen brain atlas using a custom-built MATLAB script which produced the final regional cell count.

### Anterograde and retrograde axonal tracing

Adult mice (range P31-P70) from each IT line were anesthetized with 1.5-2% isofluorane and respiration and temperature were continuously monitored throughout the procedure. Before surgery, ketoprofen (5mg/kg) and dexamethasone (0.5mg/kg) were administered subcutaneously, and lidocaine (2-4mg/kg) was applied intra-incisionally. Mice were mounted in a stereotactic frame (RWD Instruments, Shenzhen, China), and target coordinates were identified. Following a scalp incision, a craniotomy was performed using a dental drill. Pulled-glass pipettes with a 20-30μm tip diameter were backfilled with mineral oil and front loaded with viral solution. Viral injections were delivered using a NanoLiter 2010 injector (World Precision Instruments, Sarasota, FL, USA) controlled by a SMARTouch controller at a rate of 40nL/min with a total volume of 200-250nL per site.

For anterograde tracing, Cre- or Flp-dependent fluorescent tracers were packaged in an AAV9 capsid. AAVs expressing a conditional green fluorescent tracer were injected into coordinates centered over the left primary forelimb motor cortex (AP +0.7mm, ML 1.85mm, DV −0.5mm), and AAVs expressing a conditional red fluorescent tracer were injected into the right primary sensory cortex (AP +0.23mm, ML 2.62mm, DV −0.5mm). For retrograde tracing, 50nL of AAVretro expressing a Flp- or Cre-dependent green fluorescent tracer was injected into the right PERI/ECT (AP −2.65, ML −4.3, DV −3.85). After stereoinjection, the pipette was left in place for ten minutes and then slowly withdrawn at a rate of 50μm/min. The scalp was closed with surgical adhesive and mice were monitored for post-operative recovery. The incubation period was approximately 30 days before transcardial perfusion with paraformaldehyde and tissue harvesting.

| **Anterograde tracers** |  |  |
|---|---|---|
|  | **Motor cortex injection** | **Sensory cortex injection** |
| Flp | AAV9 pAAV-Ef1a-fDIO-EYFP (Addgene #55641-AAV9) | AAV9 pAAV-Ef1a-fDIO-mCherry (Addgene #114471-AAV9) |
| Cre | AAV9 pCAG-FLEX-EGFP-WPRE (Addgene #51502-AAV9) | AAV9 pAAV-FLEX-tdTomato (Addgene #28306-AAV9) |
| **Retrograde tracers** |  |  |
|  | **PERI-ECT-TEa injection** |  |
| Flp | AAVretro-EF1a-fDIO-EYFP (Addgene #55641) |  |
| Cre | AAVretro-CAG-DIO-EYFP (Addgene #104052) |  |

### Tamoxifen induction

To activate Cre-dependent axonal tracers or GCaMP in CreER IT lines, Cre recombination was induced by intraperitoneal administration of 100-200mg/kg tamoxifen (20mg/mL in corn oil) in two divided doses separated by at least 48 hours within the first five post-operative days. For STPT imaging of CreER IT lines crossed with Cre-dependent reporter lines, 100-200mg/kg tamoxifen (20mg/mL in corn oil) was injected intraperitoneally at adult ages, with a second tamoxifen dose administered seven days later. Brains were harvested at least seven days following the second tamoxifen dose.

### In-situ hybridization

Hybridization chain reaction (HCR) in situ hybridization (Molecular Instruments, Los Angeles, CA, USA) was performed according to the manufacturer’s protocol. Briefly, 40µm cryosectioned coronal brain slices were fixed in 4% PFA for five minutes, washed twice with 1X PBS, and permeabilized overnight in 70% ethanol. Sections were then rinsed twice with 1X PBS, transferred to 5% sodium dodecyl sulfate (SDS) solution, and then rinsed four times in 2X saline-sodium citrate buffer (SSC). Prehybridization was then performed with Hybridization Buffer (Molecular Instruments, Los Angeles, CA, USA) for a minimum of 30 minutes prior to application of a mixture of 2.5µL probe mix in 500µL of Hybridization Buffer. After overnight incubation at 37°C, sections were washed four times in Wash Buffer (Molecular Instruments, Los Angeles, CA, USA) at 37°C, rinsed twice in 2X SSC, and then Amplification Buffer (Molecular Instruments, Los Angeles, CA, USA) was applied at room temperature for 30 minutes. Fluorescent amplification probes were heated for 90 seconds at 90°C then snap cooled to room temperature, mixed in Amplification Buffer, and applied to the sections for overnight incubation at room temperature. Sections were then washed twice in 2X SSC, mounted on glass slides with DAPI-containing mounting medium, and imaged with a Zeiss LSM 900 confocal microscope (Zeiss, Oberkochen, Germany). Images were then processed with Zeiss ZEN software and ImageJ.

### Immunohistochemistry, microscopy, and image processing

To analyze anterograde projection patterns, every tenth consecutive 40µm coronal section from preserved brains was selected for immunostaining and confocal imaging. Sections were washed three times in 1X PBS and then fluorescent signal was amplified via immunostaining with either chicken anti-GFP (1:1000, AvesLabs GFP-1020, Aves Labs, Davis, CA, USA) or rabbit anti-RFP antibody (1:1000, Rockland 600-401-379, Rockland Immunochemicals, Limerick, PA, USA) in blocking solution (1X PBS with 1% bovine serum albumin and 0.1% Triton-X100). Sections with primary antibodies were maintained at 4°C overnight. Primary antibodies were then removed via three washes in 1X PBS and secondary antibodies – goat anti-chicken-488 (1:500, Invitrogen A11039, Thermo Fisher Scientific, Waltham, MA, USA) or goat anti-rabbit-594 (1:500, Invitrogen, A11012, Thermo Fisher Scientific, Waltham, MA, USA), as appropriate – were applied in blocking solution for one hour at room temperature before a final three washes in 1X PBS and mounting on glass slides.

Sections were first imaged at 2x magnification using a Keyence BZ-X810 inverted fluorescence microscope (Keyence, Osaka, Japan). Select sections were re-imaged at higher resolution via confocal microscopy. To do so, tiled images at 5x magnification were acquired using a Zeiss LSM 900 confocal microscope with Zeiss ZEN software (Zeiss, Oberkochen, Germany), with laser power and detector settings kept constant across all sections from one brain. All coronal sections were registered to the Allen Institute Common Coordinate Framework (CCFv3) using the Allen Brain Atlas Alignment (ABBA) pipeline in ImageJ. Initial coarse registration was performed with DeepSlice, followed by refinement using affine and spline transformations. Registrations were visually inspected for accuracy.

### Evaluating IT marker expression in mouse motor cortex in the Allen Institute single-cell transcriptomic atlas

The raw 10X single-cell RNA sequencing dataset and associated metadata for mouse whole brain and hippocampus were downloaded from the Allen Institute (https://brain-map.org/our-research/cell-types-taxonomies/cell-types-database-rna-seq-data/mouse-whole-cortex-and-hippocampus-10x; accessed November 21, 2022; Yao et al., 2023). Using the Seurat package (v5.0.0) in R, the raw data was normalized and scaled using SCTransform, principal component analysis was performed using RunPCA, and a UMAP was generated from the first 30 principal components using RunUMAP. Using the existing cell annotations provided by the Allen Institute, the dataset was subsetted to include only excitatory neurons with the following subclass labels: L2/3 IT CTX, L4/5 IT CTX, L5 IT CTX, L5 PT CTX, L5/6 NP CTX, L6 CT CTX, L6 IT CTX. Expression of individual IT markers was then assessed by further filtering excitatory neurons for those with a raw transcript count >100. A threshold of 100 adequately recapitulated the expected distribution based on Allen in situ hybridization images for each marker, our own HCR in situ hybridization data, and our reporter-crossed mouse line imaging. From the subset of cells that met this expression threshold for a given IT marker, the percent of each IT subclass label, as annotated in the Allen metadata, was calculated and graphed as a piechart using ggplot2 in R.

### IT marker co-expression analysis via MERFISH

A custom 398-gene MERFISH panel was designed using gpsFISH (Zhang et al., 2024) with the Allen Institute isocortex and hippocampal formation scRNA-seq dataset (Yao et al., 2023) as reference. The panel combined manually curated literature markers with computationally selected genes optimized for glutamatergic subclass and supertype classification, including probes for IT markers, and all genes were verified against Allen Institute in situ hybridization data (Lein et al., 2007; https://mouse.brain-map.org).

Brain samples were processed according to the manufacturer’s protocol, followed by hybridization with our custom MERFISH panel. Following image acquisition, transcript decoding and cell segmentation were performed using the Vizgen Post-processing Tool (VPT) and Cellpose. Cells with fewer than 40 or more than 3,000 transcripts, or with cell volume below 100 µm^3^ or above 3,000 µm^3^, were excluded from downstream analyses. For each marker gene, an expression-positive threshold was defined by comparing MERFISH transcript counts against the spatial expression patterns from the Allen Mouse Brain ISH dataset (Lein et al., 2007; https://mouse.brain-map.org) and our STPT data from each reporter-expressing IT line and then selecting thresholds that recapitulated the expected spatial distribution of each marker. Cells within the cortical region of interest (SSp) were then classified for each gene pair as positive for one gene only, positive for both genes, or negative for both, and plotted in spatial coordinates to visualize the distribution of expression patterns across the tissue. For each gene pair, the proportion of cells positive for one or both markers (relative to the total number of cells positive for both markers) was calculated and displayed as stacked bar plots.

### Cranial window implantation and Flp-dependent GCaMP viral injection surgeries

Cranial window surgery was performed as previously described (Li et al., 2026). Briefly, for two-photon imaging, a 3mm craniotomy was made, centered at the right SSp (AP 0.23, ML −2.62). In *Wfs1-Flp* animals, a Flp-dependent GCaMP8f virus (AAVDJ-CAG-fDIO-GCaMP8f) designed in-house was injected into the right SSp (AP 0.23 mm, ML −2.62 mm, DV −0.4 mm). No viruses were injected for *Deptor-CreER;GCaMP8m* mice. A two-layer 3mm cranial window was then implanted. For head-restrained experiments, a custom designed titanium flat headpost was secured on the skull with dental cement (C&B Metabond, Parkell, Edgewood, NY, USA or Ortho-Jet, Lang Dental, Wheeling, IL, USA). *Deptor-CreER;GCaMP8m* mice were induced with tamoxifen post-operatively to activate Cre-dependent GCaMP expression, as described above.

### Reach-withdraw-drink (RWD) behavior paradigm

Details of the RWD task have been described previously (Galinanes et al., 2018; Li et al., 2026). Briefly, animals were trained in the dark with their head fixed and right arm blocked. Mice were trained to grasp sucrose solution drops (10%, 20-50μL) delivered in the peri-orofacial space, with the waterspout initially placed to the left of the mouse’s snout. Training consisted of one session per day for three days, with 150 trials per session and random inter-trial intervals of 12-20 seconds. Once mice consistently performed RWD towards the left location (hit rate > 95%), the spout would then be programmed to randomly move to one of three equidistant locations (identified as left P1, center P2, and right P3 relative to the snout). Water was delivered after a random delay (3-7 seconds) via a solenoid valve. The mouse was given sufficient time to finish RWD (7+ seconds) before the trial ended.

### Two-photon imaging of calcium-dependent activity

Single-neuron activity was recorded in awake, behaving mice via two-photon microscopy (Ultima Investigator, Bruker, Billerica, MA, USA) using a fs-pulse Ti:Sapphire laser (Mai Tai DeepSee, Spectra-Physics, Milpitas, CA, USA) at 920nm excitation. GCaMP signals were isolated with a 525/70nm bandpass filter and collected using gallium arsenide phosphide (GaAsP) photomultiplier tubes (PMTs). Imaging was conducted in single-plane mode at 512×512 pixel resolution and ∼30 Hz frame rate, with each trial spanning 475 consecutive frames. Source laser power was set between 150–400mW, corresponding to 33–76mW at the objective.

### High speed behavior tracking

Reaching behavior was captured from the front and left side of the animal using high-speed cameras (Point Grey Flea3, Teledyne Vision Solutions, Thousand Oaks, CA, USA) at 150 fps. Camera geometry was calibrated using the Camera Calibrator app in MATLAB, and videos were acquired and saved via custom MATLAB code for offline analysis. Body part trajectories were tracked using two DeepLabCut (Mathis et al., 2018) models trained separately for each camera view, yielding 18 and 22 labeled landmarks in the front and side views, respectively, including digits, facial features (nose, mouth, tongue, eye), the waterspout, and water droplet. Detailed characterization of the behavior was described in previous publications (Li et al., 2026). All data processing and analyses were carried out in MATLAB or Python unless otherwise noted.

### Two-photon imaging analysis

Motion correction and region of interest (ROI) extraction were performed using the Suite2P package (Pachitariu, 2017). ROIs were retained for further analysis if they met the following criteria: classifier probability > 0.05, radius > 6 pixels, and skewness > 0.7. Neuropil contamination was corrected as *F*_neuropil_corrected_(*t*) = *F*(*t*) – 0.7 x *F*_neuropil_(*t*), where F represents raw fluorescence extracted from ROI centroids. Fluorescence changes were calculated as ΔF/F₀, with F₀ defined as the mean baseline fluorescence over the one second window immediately preceding water dispense.

To examine the relationship between neuronal activity and behavior, calcium traces were aligned to discrete behavioral events: water onset (valve trigger signal), lift onset (DeepLabCut), spout touch (DeepLabCut), and hand lick (DeepLabCut). All behavior experiments were conducted with randomization. Sample sizes were determined based on established precedents in the literature (Li et al., 2026), with sufficient trials collected per condition to ensure robust and reproducible results. Both male and female animals were included to avoid sex bias.

## ACKNOWLEDGEMENTS

The authors would like to thank Myra Han for assistance with mouse husbandry and tissue processing. The authors would also like to thank Mariana Holguin Lopez for assistance with MERFISH processing.

## Supplementary Figure

**Supplementary Figure 1: Demonstration of IT marker expression patterns across multi-modal mouse brain atlases. (A)** Representative sagittal cortical images of in situ hybridization (ISH) for each IT marker demonstrating different laminar patterns of expression. Images obtained from the Allen Institute ISH Mouse Brain Atlas (https://mouse.brain-map.org/). **(B)** Reanalysis of single-cell gene expression data from whole mouse brain (Yao et al., 2023) from the Allen Institute (https://brain-map.org/our-research/cell-types-taxonomies/cell-types-database-rna-seq-data/mouse-whole-cortex-and-hippocampus-10x). Only cells annotated by the Allen Institute as having an IT class identity are included in the Uniform Manifold Approximation and Projection (UMAP; right) with original Allen Institute laminar subclass annotations (L2/3 IT, L4/5 IT, L5 IT, and L6 IT). Feature plots for individual IT markers (left) demonstrate different densities and intensities of relative gene expression across IT subclasses (purple indicates higher relative expression; gray indicates no expression). **(C)** Pie charts depicting the percent of each IT subclass (as originally annotated by the Allen Institute) within the total population of marker-expressing excitatory cortical neurons. Scale bars = 250 µm.

**Supplementary Figure 2: Spatial expression and co-expression of IT markers. (A)** Representative HCR ISH images of *Wfs1* and *Igfn1* co-expression in layer 2/3 and *Ptprm* and *Deptor* co-expression in layer 5. Scale bar = 50 µm. **(B)** Representative coronal brain cross-section from a wild type mouse processed for MERFISH. Cyan wedge indicates sampled region of SSp from which expression data in D-F were generated. **(C)** Color code for marker-positive cells assessed via MERFISH in D-F. **(D)** MERFISH data illustrating the co-expression (pink) of each IT marker with the pan-IT marker *Satb2* (dark gray) along with corresponding quantitative bar graphs. **(E)** MERFISH data illustrating the co-expression (pink) of upper-layer IT markers. **(F)** MERFISH data illustrating the co-expression (pink) of lower-layer IT markers. For this illustration, MERFISH was performed in N=1 cortical sections from N=1 mice.

**Supplementary Figure 3: Subcortical expression patterns of upper-layer IT markers. (A)** The top 20 subcortical regions with the highest densities of *Cux1*^+^ cell bodies are plotted in descending order. **(B)** Representative examples of subcortical regions with dense *Cux1*^+^ cell bodies from *Cux1-Flp;Ai65F* mice are shown. **(C)** The top 20 subcortical regions with high densities of *Wfs1*^+^ cell bodies are plotted. **(D)** Representative images of dense *Wfs1*^+^ cell bodies in subcortical regions of *Wfs1-Flp;Ai65F* mice. **(E)** The top 20 subcortical regions with high densities of *Igfn1*^+^ cell bodies are plotted. **(F)** Representative images of piriform cortex and retrosplenial cortex with dense *Igfn1*^+^ cell bodies in *Igfn1-Flp;Ai65F* mice are shown. Cell count data are derived from N=3 mice for each mouse line and mean cell count per region is plotted. Error bars represent standard deviation. Scale bars in A = 500 µm. All other scale bars = 1000 µm. ACB = nucleus accumbans, AI = agranular insular cortex, AON = anterior olfactory nucleus, CA1 = CA1 of the hippocampus, CLA = claustrum, ENTl = lateral entorhinal cortex, FS = fundus of the striatum, IA = intercalated nucleus of the amygdala, LA = lateral amygdalar nucleus, MEA = medial amygdalar nucleus, MOB = main olfactory bulb, OT = olfactory tubercle, PAR = parasubiculum, PERI = perirhinal cortex, PIR = piriform cortex, RSP = retrosplenial area, SSp = primary somatosensory area, VISpor = postrhinal area

**Supplementary Figure 4: Subcortical expression patterns of lower-layer IT markers. (A)** The top 20 subcortical regions with the highest densities of *Kcnh5*^+^ cell bodies are plotted in descending order. **(B)** Representative examples of subcortical regions with dense *Kcnh5*^+^ cell bodies in *Kcnh5-CreER;Ai14* mice are shown. **(C)** Top 20 densest subcortical regions for *Deptor*^+^ cell bodies are plotted. **(D)** Representative images of subcortical regions with dense *Deptor*^+^ cell body labeling in *Deptor-CreER;Ai14* mice. **(E)** Top 20 densest subcortical regions for *Ptprm*^+^ cell bodies are plotted. **(F)** Representative images of subcortical regions with dense *Ptprm*^+^ cell body labeling in *Ptprm-CreER;Ai14* mice. Cell count data are derived from N=3 mice for each mouse line and mean cell count per region is plotted. Error bars represent standard deviation. Top middle scale bar in (B) and middle scale bar in (C) = 1000 µm, all other scale bars = 500 µm. AON = anterior olfactory nucleus, ARH = arcuate hypothalamic nucleus, BAC = bed nucleus of the anterior commissure, BST = bed nuclei of the stria terminalis, chpl = choroid plexus, CP.vl = caudoputamen ventrolateral, IA = intercalated amygdalar nucleus, LA = lateral amygdalar nucleus, MEA = medial amygdalar nucleus, MOB = main olfactory bulb, OT = olfactory tubercle, PA = posterior amygdalar nucleus, PG = pontine gray, PIR = piriform area, POST = postsubiculum, PRE = presubliculum, PVT = paraventricular nucleus of the thalamus, SUB = subiculum, TMv = tuberomammillary nucleus ventral part, VMH = ventromedial hypothalamic nucleus

