## Supplementary figures and images for "An expanded toolbox for genetic access to molecularly defined cortical intratelencephalic neuron subtypes"

### Supplementary Figure 1

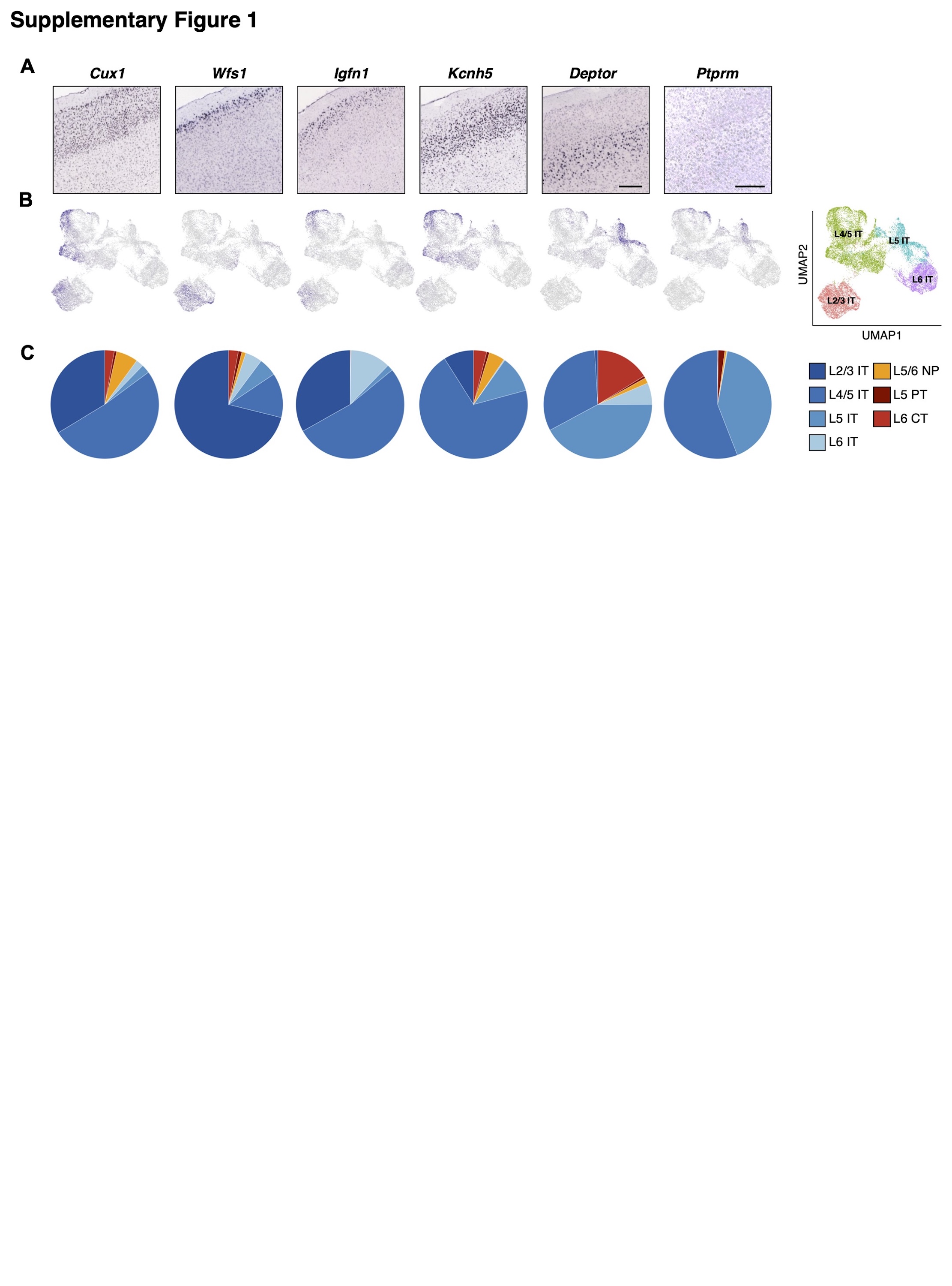

### Supplementary Figure 2

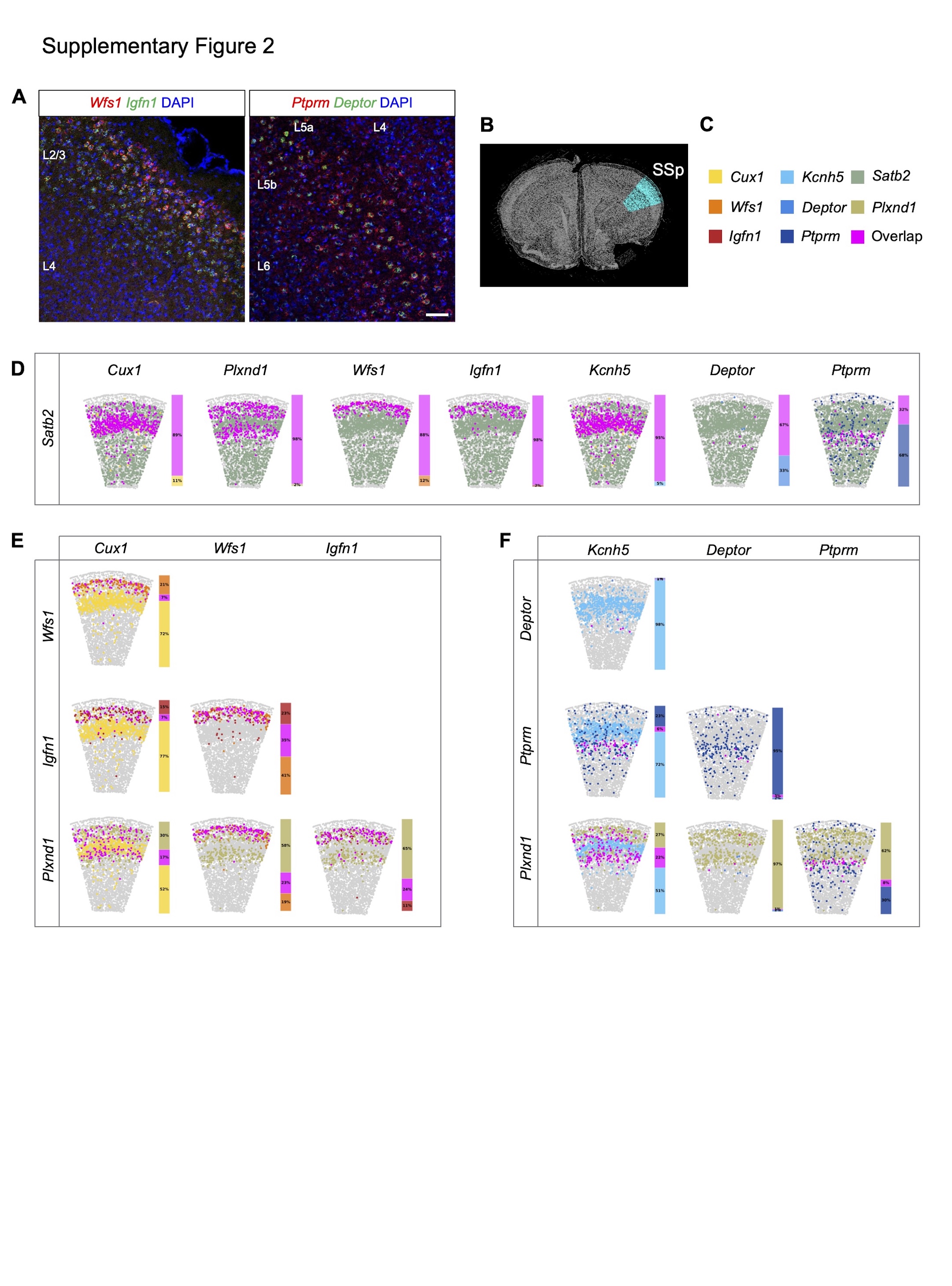

### Supplementary Figure 3

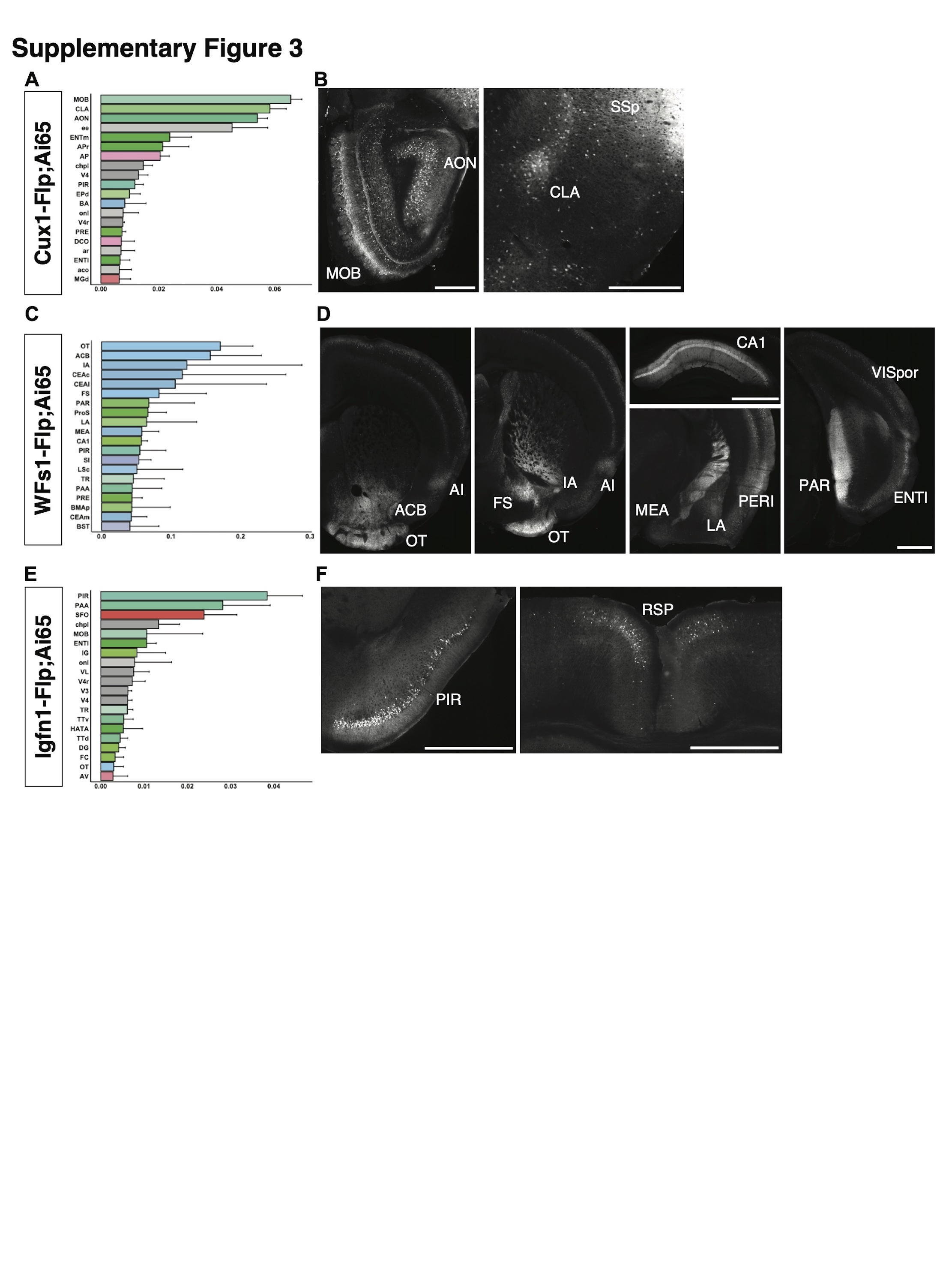

### Supplementary Figure 4

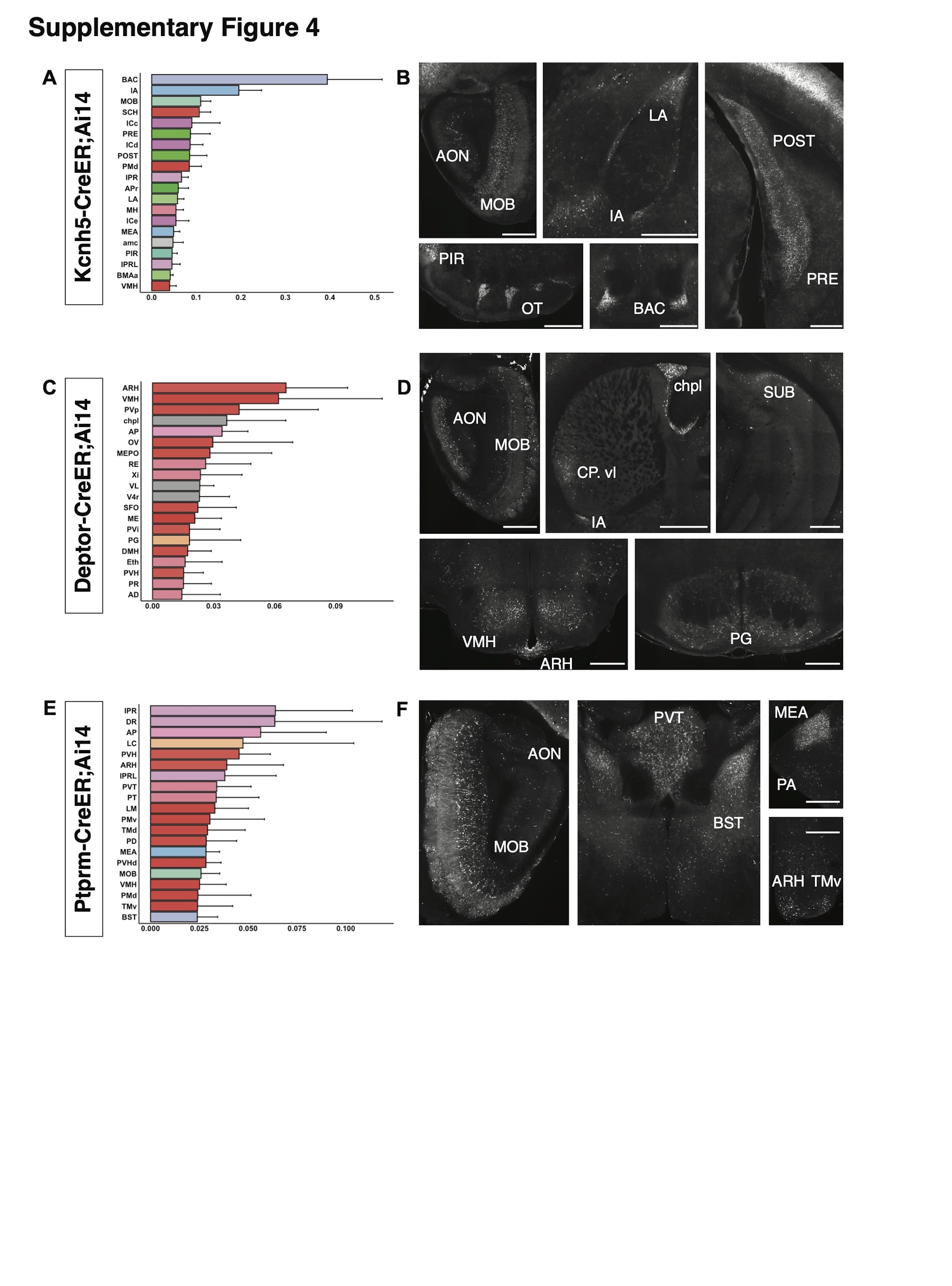
